# Learning the sound inventory of a complex vocal skill via an intrinsic reward

**DOI:** 10.1101/2022.09.27.509691

**Authors:** Hazem Toutounji, Anja T. Zai, Ofer Tchernichovski, Richard H.R. Hahnloser, Dina Lipkind

## Abstract

Reinforcement learning (RL) is thought to underlie the acquisition of vocal skills like birdsong and speech, where sounding like one’s “tutor” is rewarding. Yet, we find that the standard actor-critic RL model of birdsong learning fails to account for juvenile zebra finches’ efficient learning of an inventory of multiple syllables. But when we replace a single actor with multiple independent actors that jointly maximize a common intrinsic reward, then birds’ empirical learning trajectories are accurately reproduced. Importantly, the influence of each actor (syllable) on the magnitude of global reward is competitively determined by its acoustic similarity to target syllables. This leads to each actor matching the target it is closest to, and occasionally, to the competitive exclusion of an actor from the learning process (i.e., the learned song). We propose that a competitive-cooperative multi-actor (MARL) algorithm is key for the efficient learning of the action inventory of a complex skill.

## Introduction

Animal behavior provides a unique opportunity for understanding evolutionary solutions to complex learning problems. One prime example is learning the inventory of components for combinatorial vocal skills such as speech sounds or birdsong syllables. In both humans and songbirds, the acquisition of vocal skills is thought to be subserved by a reinforcement learning (RL) mechanism^1–3^, as evidenced by dopamine signaling ^4–8^. Dopaminergic neurons signal reward prediction error (RPE) by increasing or decreasing their firing rate when an appetitive outcome is respectively better or worse than expected^9–12^. Similarly, in adult songbirds, dopaminergic projections to the basal ganglia signal whether aversive singing outcomes imposed by distorted auditory feedback are better or worse than expected^8^. Such error signal coding is at the heart of many hypothesized RL mechanisms of developmental birdsong learning^7,13,14^.

The goal of reinforcement learning is to maximize rewards, but internally motivated processes such as speech or birdsong learning readily occur in the absence of external rewards. Learning must therefore rely on intrinsically generated reward signals that are contingent on the similarity between current and target performance. It is not known what kind of RL mechanism, and which form of intrinsic reward drive the acquisition of an inventory of sounds necessary for performing combinatorial vocal sequences.

Here we attempt to infer the intrinsic reward underlying the learning of a syllable inventory in zebra finches. We assess inventory learning in terms of pitch, which zebra finches readily imitate^15,16^. Zebra finches possess vocal combinatorial ability^17^, despite typically singing a fixed syllable sequence. Juveniles learn the syllable inventory of their target song (a memorized song of an adult^18^) independently of syllable order, using a highly efficient “greedy” strategy^16^. Namely, they make the minimal necessary changes to the sounds in their own vocal repertoire to match the syllables in the target song. This suggests a dedicated reward computation for syllable inventory learning that is not contingent on sequential order. Our goal is to determine the functional form of the reward – namely, how it is contingent on the pitch similarity between the syllables a bird performs and the syllables in its target song.

We develop a multi-actor reinforcement learning (MARL) model where independent RL actors control the performance of distinct syllables. Akin to multi-agent learning systems^19^, actors maximize a common intrinsic reward by nullifying an RPE^8^. We implement the greedy and order-independent learning observed in zebra finches by competitive-cooperative interactions among actors: 1. actors compete over target syllables leading to each target being matched by the most similar actor; 2. actors cooperate to maximize reward that increases in proportion to the number of matched targets. Such interactions require a reward function that takes all possible pairwise actor-target comparisons into account.

We test our model against several computationally simpler alternatives on the task of simulating the empirical learning trajectories ^16^ of juvenile male zebra finches that are experimentally induced to learn new syllables. We find excellent agreement with data for a competitive-cooperative reward model that acts over short distances (between actors and targets) in pitch space. The model successfully predicts a competitive hierarchy between syllables and calls in an experimental test where we incite birds to exclude an existing syllable from a song and replace it with a call. We conclude by presenting the predictions of our model for the responses of dopaminergic projections to the songbird basal ganglia during the learning of a vocal inventory.

## Results

### Vocal imitation of a single syllable best agrees with a light-tailed distribution of intrinsic reward

We started with a simple one-syllable learning problem – a juvenile bird learns to adjust the pitch of a single syllable to resemble a target syllable sung by an adult “tutor” (Figure 1A). We modeled the learning system as an RL agent consisting of an actor and a critic. This agent attempts to maximize an intrinsically generated reward *R* that is inversely related to pitch difference Δ between the bird’s syllable and the target. The actor is a motor program that samples variable syllable instances from an underlying *performance distribution* parameterized by its mean *S* (Figure 1B). After each instance, an intrinsic reward *R* is generated from a *reward distribution* that is centered on a perfect imitation of the target syllable *T*. The actor is associated with a critic that expects maximal reward for renditions near the actor’s mean performance; the critic computes the expected reward or quality *Q* from a distribution with the same functional form as the reward but centered at *S*. The actor learns by computing the difference between the received reward *R* and the quality *Q*. This difference *δ* = *R* − *Q* is the RPE thought to be encoded by dopaminergic neurons^8,20^. Learning is driven by minimization of the square RPE (Figure 1C). Each syllable instance leads to an update Δ*S* of the actor’s mean performance *S* according to a simple iterative rule (equation 7; Materials and Methods). This update brings the quality *Q* of the instance closer to the actual reward *R* and makes *S* more similar to the target *T* (see Materials and Methods). Syllable instances in the vicinity of *S* almost always result in a negative RPE. That is because the critic expects maximal reward near *S*, whereas the actual reward is maximal only when *S* matches *T*. However, the RPE is larger (less negative) for instances on the target side of *S* than instances on the opposite side, which is what shifts *S* towards *T*. By construction, the RPE is zero and learning stops when *S* coincides with *T*.

**Figure 1:**
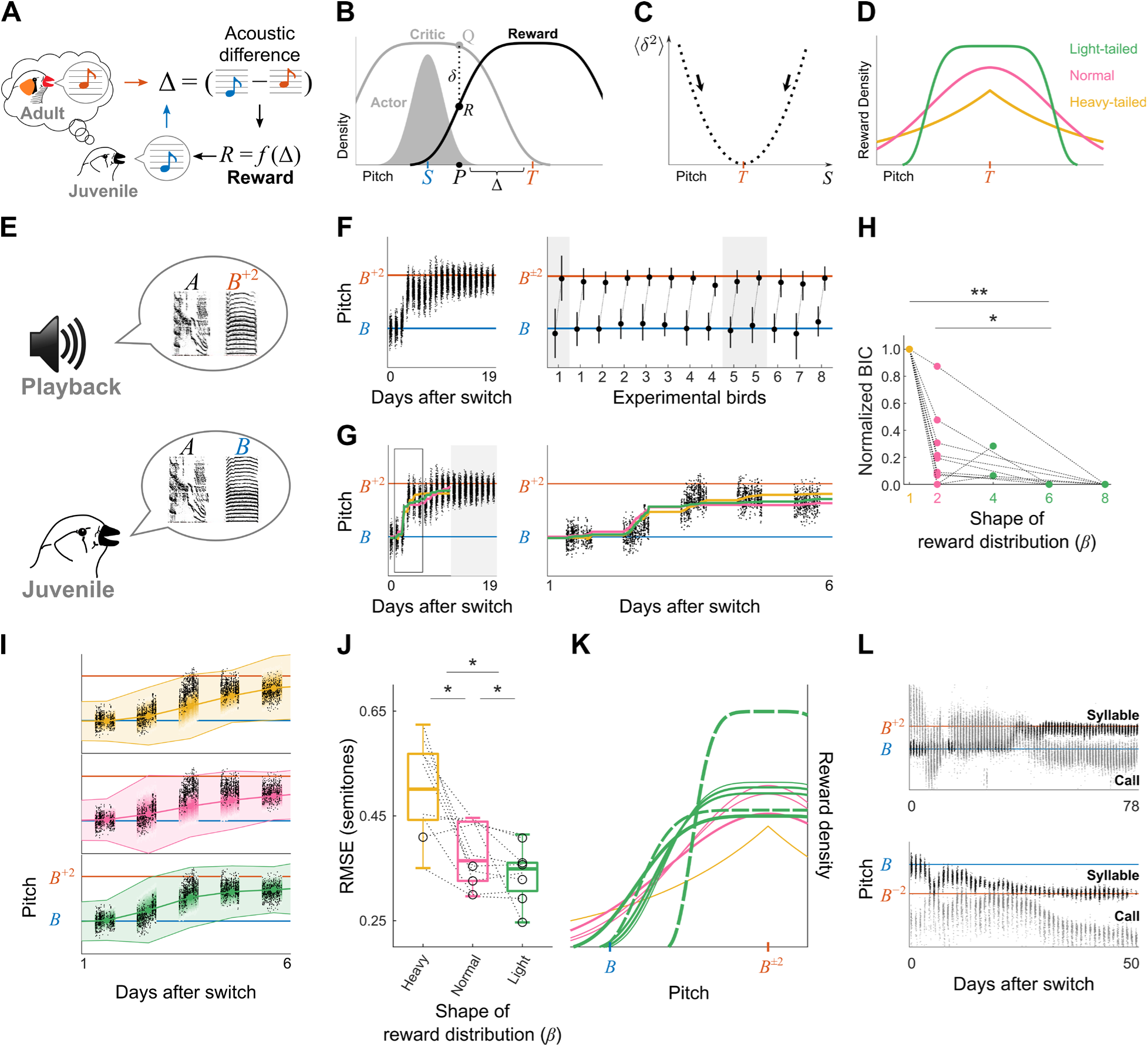
Vocal imitation of a single syllable best agrees with a light-tailed distribution of intrinsic reward. **A**. A juvenile male zebra finch adjusts the performance of a syllable (blue) to acoustically match its target – an adult “tutor’s” syllable (orange); learning maximizes an intrinsic reward *R* that is inversely related to pitch difference Δ between the juvenile’s syllable and the target. **B**. An actor-critic model for learning a syllable: the actor is a motor program that generates syllable instances *P* from a distribution with mean *S*. The critic estimates an expected reward *Q* for each instance and compares that to the intrinsic reward *R*, resulting in a reward prediction error (RPE) *δ* = *R* − *Q*. We assume that *Q* and *R* have identical distributions, with *R* being maximal at a perfect imitation of the target (Δ = 0), and *Q* being maximal at the motor program mean *S*. **C.** The motor program mean *S* is updated to minimize the mean squared RPE *δ*, which is at its minimum when *S* equals *T* (see Eqs. 6 and 7 in Materials and Methods). **D.** Hypothetical intrinsic reward distributions modelled as a generalized normal distribution. The shape parameter beta determines whether a distribution is light-tailed (*β* > 2; green), heavy-tailed (*β* < 2; yellow) or normal (*β* = 2; magenta). **E.** A syllable-matching experiment: a bird that has learned a two-syllable song *AB* is induced to learn a new song, *AB*^±2^, where the pitch of syllable *B*^±2^ is two semitones above or below that of *B* (an *AB* → *AB*^+2^ experiment is shown). **F**. Left, learning trajectory of an experimental bird trained with the task in **E**, showing the median pitch of consecutive instances of syllable *B* after the presentation of the new tutor song (day 0); blue and orange lines represent the pitch of syllables *B* and *B*^+2^, respectively. Right, learning outcomes in all experimental birds trained with 2-semitone pitch shifting tasks, showing pitch distributions (median and 95% confidence interval, CI) at the start point and end point of learning for each shifted syllable, connected by dotted lines (13 syllables in n = 8 birds; orange line, the absolute pitch difference between *B* and *B*^±2^). Grey shading denotes 3 syllables excluded from parameter estimation due to lack of convergence (see Figure S1A). **G**. Left, inferred mean pitch shifting trajectories fitted with MLE to the learning trajectory of the experimental bird in **F** left, assuming reward distributions with different values of *β* (colors as in **D**). Right, zoom-in on the learning part of the trajectory (see Figure S1B). **H**. Bayesian Information Criterion (BIC) for each syllable trajectory, assuming reward distributions with different shapes (values of *β*; dashed lines connect same-syllable data; n = 10). BIC scores for each syllable are normalized such that 1 corresponds to the worst and 0 to the best fit. Light-tailed reward improves BIC scores compared to heavy-tailed and normal reward (*=p<0.05, **=p<0.01; Benjamini-Hochberg corrected Wilcoxon signed-rank test; cf. Figure S2). **I.** Observed (black) and bootstrapped (color) pitch trajectories of the pitch-shifted syllable shown in **F** left. Shaded area represents median ± 50% CI of simulated trajectories. Colors as in **D**. Color gradient represents bootstrap density. **J.** Root-mean-square error (RMSE) between observed and bootstrapped learning trajectories (dashed lines connect same-syllable data; open circles represent the RMSE of the best reward model per syllable). Light-tailed reward improves goodness-of-fit compared to heavy-tailed and normal reward (p < 0.05, Benjamini-Hochberg corrected Wilcoxon signed-rank test; see Figure S2). **K.** Best (smallest RMSE) reward distribution for each empirical learning trajectory (colors as in **D**; solid green: *β* = 4, dashed green: *β* = 6). Thicker lines represent larger improvement by the best model compared to the second best. **L.** Two examples of a syllable that shifted to match a target although a call was initially closer.

We estimated the shape of the performance distributions of zebra finch syllables from published data^16^, and found them to be Gaussian (see Materials and Methods). To estimate the distribution of the putative intrinsic reward *R* as a function of pitch difference Δ, we examined a continuum of hypothetical reward distributions *R*(Δ), which we modeled as a generalized normal distribution^24^ with two unknown parameters: the shape *β* and scale *ς* (Figure 1D; see Materials and Methods). The scale parameter *ς* determines the standard deviation, and consequently the width of the distribution. The shape parameter *β* produces a normal distribution for *β* = 2 as a special case. Smaller values (*β* < 2) correspond to heavy-tailed distributions, and larger values (*β* > 2) correspond to light-tailed distributions. Heavy-tailed reward distributions exert long-range influence over syllables at a large pitch difference from the target. Light-tailed distributions exert short-range influence, rewarding only syllables that are relatively similar to the target. While previous studies have proposed either short^21^ or infinite range^3^ of differences from a target that can affect syllable performance, the actual range is unknown.

We inferred the model parameters *β* and *ς* by fitting our actor-critic model to actual learning trajectories from syllable-matching experiments (part of which were previously published^16^). Juvenile males were trained with artificial song tutors^16^ to shift the pitch of a syllable by two semitones to match a syllable in a tutor’s song (Figure 1E), or to shift two syllables towards two different targets (in the latter case, we treated the syllables’ learning trajectories as independent of each other). This task was accurately accomplished (Figure 1F) regardless of whether the pitch was shifted up or down, which agrees with our assumption that the intrinsic reward distribution is symmetrical (13 syllables in 8 birds; 3 syllables were excluded from further analysis due to oscillating trajectories; Figure S1)^16^. Because we were interested in whether reward functions are qualitatively sub- or super-Gaussian, we evaluated fit quality as a function of *β* by fixing *β* to distinct values and inferring *ς* via maximum likelihood estimation, or MLE (Figures 1G and S1B; see Materials and Methods).

In 8 out of 10 trajectories on which model parameters were inferred, light-tailed reward distributions (*β* = 4, 6, or 8) resulted in better fits (i.e., lower Bayesian Information Criterion, or BIC, scores) than normal (*β* = 2) and heavy-tailed (*β* = 1) distributions (Benjamini-Hochberg corrected Wilcoxon signed-rank test on non-normalized BICs; Figures 1H and S1B). We obtained similar results when we estimated the *β* jointly with other model parameters (Figure S2A). BIC scores alone, however, are insensitive to overfitted models that fail to generalize to unseen data. Therefore, we also performed a bootstrap analysis by simulating multiple learning trajectories for each ML estimate of *β* (Figures 1I and S1B). Light-tailed reward distributions resulted in improved goodness-of-fit (i.e., smaller Root-Mean-Square Error, or RMSE), between empirical and simulated trajectories in 6 out of 10 cases (Figures 1J and S2B); in 4 of 10 cases in which the light-tailed model was not the best, the decrease in goodness-of-fit was small compared to heavy-tailed and normal models (Figure 1K). These results suggest that a light-tailed reward distribution guides the shift in syllable performance towards a match with a target. Namely, a syllable instance triggers a non-zero intrinsic reward only if it is sufficiently similar to a target syllable. This solution is a special case of the multi-syllable inventory learning problem that we turn our attention to next.

### Imitation of a multi-syllable inventory best agrees with light-tailed reward distributions combined in a hierarchical sum-max operation

As a first step in modeling the learning of a multi-syllable inventory, we investigated the respective roles of syllables and calls in learning. Birds use both syllables and calls in their vocal repertoire to match syllables in their target song^16,17,22^, but previous findings^16^ suggest that calls are only used in cases where there are more targets than syllables in a bird’s song. We reinspected previously published data, and indeed found that when there is an equal number of syllables and targets, birds used syllables to match targets even when there were acoustically closer calls in their repertoire (figure 1L). We therefore decided to model inventory learning as a hierarchical process that prioritizes syllables over calls: a call that is similar to a target will be ignored when a less similar syllable is also present within reward range.

Next, we set up to extend our reward model for learning a single target (Figure 1B-C) to the parallel learning of multiple targets (taking into account both syllables and calls in the bird’s repertoire, Figure 2A). For that purpose, we used the inferred reward distribution contingent on the pairwise pitch difference between a syllable and a target (Figure 1G-K) as a functional component (or a basis function); we defined a global inventory reward *R* as a function of all reward components *R_ij_* associated with pairing target *i* with syllable (or call) *j* (Figure 2B). Importantly, this approach allowed us to generalize the model from single to multiple targets without introducing any new model parameters.

**Figure 2:**
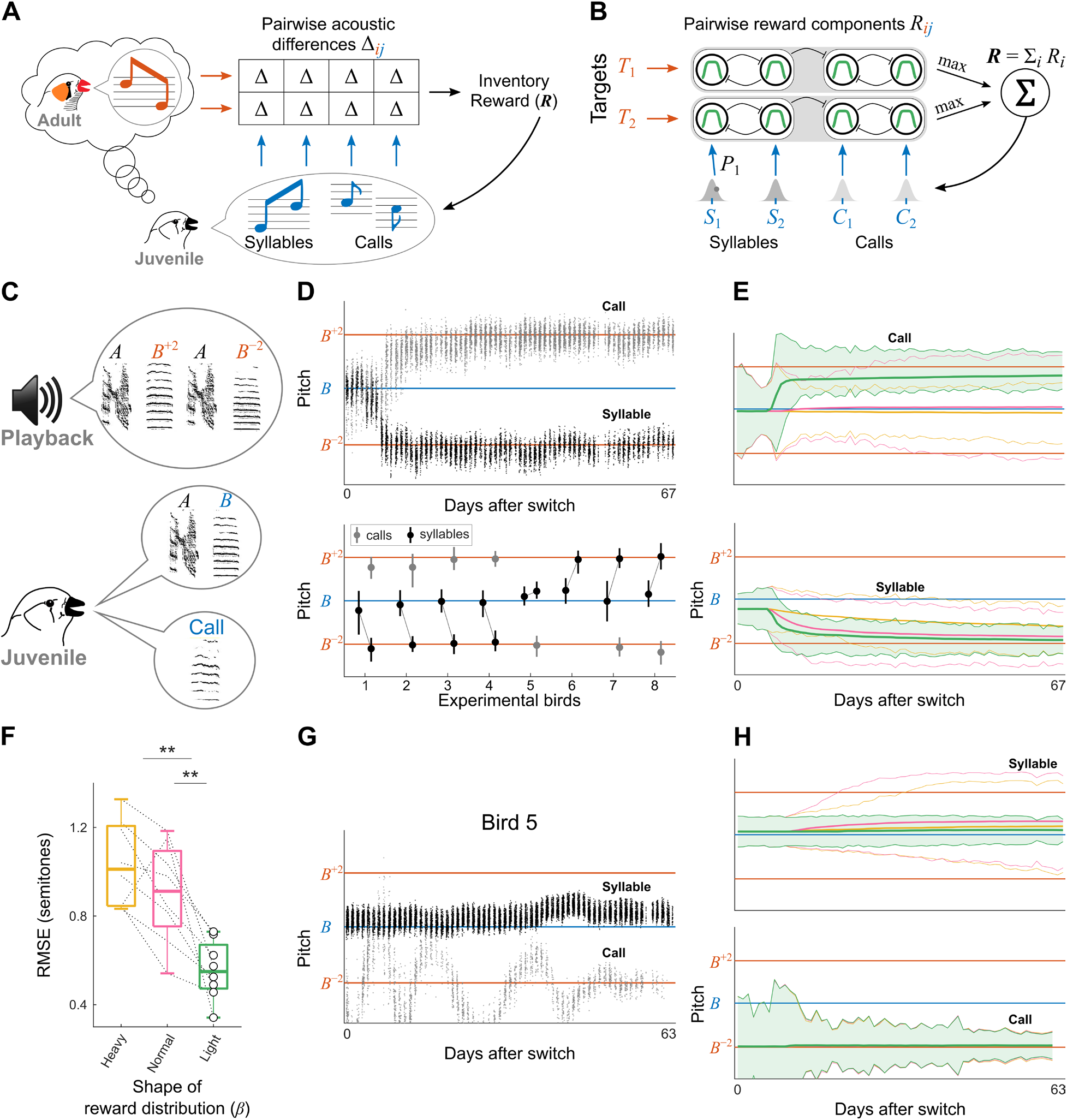
Imitation of a multi-syllable inventory best agrees with light-tailed reward distributions combined in a hierarchical sum-max operation. **A.** Learning a multi-syllable inventory, driven by a putative intrinsic scalar inventory reward *R*, which depends on the pairwise pitch differences Δ*_ij_* between the learner’s vocalizations (syllables and calls) and the targets provided by a tutor. **B.** A hypothesized computation of inventory reward *R* from pairwise light-tailed reward components *R*_*ij*_ (circles) that are contingent on syllable-target (or call-target) pitch differences. The reward ***R*** is the sum of partial rewards *R*_*i*_ computed for each target *T*_*i*_ via a hierarchical max operation over pairwise reward components. Inhibition (curved lines) among pairwise reward components leads to competition among actors and inhibition from syllables onto calls leads to prioritization of syllables over calls. **C.** Example of a multi-syllable matching experiment, in which a bird that has learned a two-syllable song *AB* is induced to learn a new song with two new target syllables *B*^+2^ and *B*^−2^, with pitch two semitones above and below that of *B*, respectively. The bird’s repertoire includes song syllables and calls. **D.** Top, pitch trajectory in one experimental bird trained with the task in **C** (notation as in Figure 1F): the bird matched one target by shifting the pitch of song syllable *B* (black), and the other target by shifting the pitch of a call (grey). Bottom, results across experimental birds (n = 8), showing pitch distributions at start point and endpoint of learning as in Figure 1E (see also Figure S4A). 7 out of 8 birds matched one target with a song syllable (black circles), and the other target with a call (grey circles). Bird 5 matched only *B*^−^ with a call but did not shift syllable *B* to *B*^+^. **E.** Bootstrapped pitch-trajectory distributions of the call (top) and syllable (bottom) of the experimental bird shown in **D** top. Median ± 50% CI are shown for the light-tailed (shaded), normal and heavy-tailed reward models (colors as in Figure 1D; also see Figure S4B). **F.** Root-mean-square error (RMSE) between experimental and bootstrapped syllable learning trajectories. Notation as in Figure 1I. Light-tailed rewards improve goodness-of-fit compared to heavy-tailed and normal rewards (p<0.01, Benjamini-Hochberg corrected Wilcoxon signed-rank test, see also Figure S4D). **G.** Pitch trajectories of song syllable *B* and call in an experimental bird that failed to match one of the targets (bird 5 in **D** bottom). **H.** Same as **E** for the bird shown in **G** (top: syllable; bottom: call). Only the light-tailed reward model predicts the failure to shift song syllable *B* towards a target.

The global reward function *R* must reproduce the context-independent and greedy syllable learning trajectories observed in zebra finches. Context-independence with respect to target positions can be achieved via the sum operation due to its associative property (a sum does not depend on the order in which the summands are added, i.e., it is permutation insensitive). Greedy matching of targets by syllables can be achieved by the maximum operation (which ignores all values smaller than the maximum). Accordingly, we constructed the inventory reward *R* as a sum-max operation over pairwise reward components *R_ij_* (Figure 2B): namely, it is a sum *R* = Σ*_i_ R_i_* of partial rewards, where the partial reward *R_i_* = *max_i_ R_ij_* associated with target *i* is given by the maximal pairwise reward that can be obtained for that target, i.e., it is computed with respect to the most acoustically similar vocalization. To implement the prioritization of syllables over calls, the maximum operation in our model is a hierarchical two-stage process: in a first stage, the maximum is taken over syllables, and if no syllable is within the reward range, then in a second stage, the maximum is taken over calls (see Materials and Methods and Figure S3A). Note that such a two-stage maximum operation can naturally emerge from a light-tailed reward component because the latter approximates a decision-making process driven by reward being either inside or outside some range. In contrast, normal and heavy-tailed models are less binary in nature and so an explicit decision-making step would be required to implement a prioritization of syllables over calls.

The inventory reward *R* is delivered to a group of actors (each as in Figure 1B), one actor per vocalization type (syllable or call; Figure 2B). Actors produce vocalizations independently of each other, and each learns to adjust its mean performance based on the identical reward *R* they receive. We assume that reward *R* is delivered after each syllable or call instance (rather than at the end of a song) and that partial rewards *R_i_* are computed with respect to the syllable or call instance just performed and the means *S* (i.e., motor memories) of the other syllables and calls in the bird’s current inventory (see Materials and Methods and Figure S3B). Upon production of an instance, an actor learns from the RPE which is defined as the difference between the inventory reward *R* and an expected reward *Q* computed by this actor’s critic (see Eq. 11 in Materials and Methods). By construction, the RPE is zero and learning stops when each target is matched by an actor’s mean.

To test our model, we used data of a published set of multi-syllable learning experiments^16^, where juvenile males were induced to learn two new targets, both of which resembled a single syllable in their current song (i.e., were pitch-shifted by two semitones up and down from that syllable, Figure 2C). Birds predominantly shifted the pitch of the syllable to match one of the new targets and recruited a call to match the other target (7 of n = 8 birds, Figures 2D and S4A). Consistent with a prioritization of syllables over calls, the song syllable shifted towards the target to which it was more similar (Figures S4A and S4B, left), while the call shifted to the other target, even when it was more similar to the target matched by the syllable (3 out of 8 cases, Figures S4A and S4B, right; these findings were not published in Lipkind et al., 2017^16^).

We compared simulated and empirical learning trajectories (as in Figure 1I), in terms of both learning endpoint and trajectory shape. We compared simulated learning trajectories generated by light-tailed, normal, and heavy-tailed variants of the model (with model parameters as inferred in Figure 1, except for the performance distributions’ initial means and daily variances which we estimated in each bird separately). The light-tailed model was significantly better at matching the empirical trajectories (Figures 2E-2F, S4C, left and middle, and S4D), including the replication of a failure of a bird to match one of the targets (Figures 2G-H). In that bird, the performance distribution of the song syllable is relatively narrow, and consequently lies outside the range of light-tailed reward distributions but is within the ranges of normal and heavy-tailed distributions. Therefore, only the light-tailed distribution correctly predicts the failed shifting of the syllable in this bird (Figure 2H, top).

The light-tailed variant of our sum-max MARL model produces good fits with empirical data, but a sum-max reward is computationally intense, requiring all possible pairwise similarity evaluations at each step. We tested two alternative reward models to see if simpler reward computations can provide comparable results. First, we tested a sequential-order-dependent (SOD) model, previously proposed by Fiete and colleagues^3^ (see Materials and Methods). In the SOD model, the reward delivered after each vocalization instance is a function of the similarity between that instance and the temporally aligned target (rather than of all possible pairwise similarities). In addition, target syllables have an infinite range of influence – i.e., they can attract even very dissimilar vocalizations. This is achieved via a binarized reward with an adaptive threshold that guarantees equal numbers of rewarded and unrewarded instances at any pitch distance from a target^3^. In contrast to the sum-max model, which made the correct (empirically observed) assignment of syllables to targets in 8 out of 8 cases, the SOD model succeeded in 3 cases only (Figure S4E, SOD 1 model). The context-dependent evaluation algorithm in the SOD model led to an incorrect assignment of temporally aligned targets in 4 cases where birds shifted a syllable to a misaligned target, while the binarized reward computation did not predict the observed failure of one bird (Figure 2G) to shift a syllable to a target. In total, the SOD model resulted in a large simulation error in 5 out of 8 birds, compared to near zero simulation error with the sum-max model. We also tested a variant of the SOD model with a light-tailed reward computation (SOD 2 model; see Materials and Methods), which correctly predicted the case of failed shifting, but did not predict the matching of misaligned targets, resulting in very large errors in 4 out of 8 birds (Figure S4E).

Next, we tested a reward model, which like the sum-max MARL model, is context-independent and competitive, but where instead of actors competing over targets, the targets are competing over actors (a sum-over-actors MARL model). Instead of selecting the most acoustically similar actor for each target, we select the most acoustically similar target for each actor, then sum the reward contributions of the winning targets, so that each actor ends up matching the target that is most similar to it. This model is computationally simpler than the sum-max model, since each step involves only the pairwise comparisons between a given actor and each of the targets, rather than all possible comparisons. However, the sum-over-actors model would not ensure that all targets are matched in cases where a target is not closest to any syllable or call. We therefore tested model predictions in two cases where the syllable and the call start-points were closer to the same target and were both within the reward range of both targets (Birds 1 and 3, Figure S4C, right). We ran simulations of the sum-over-actors model using heavy-tailed, normal, and light-tailed reward components as before. In each case, the sum-over-actors model predicted that the call and syllable will match the same target, contrary to the empirical data. In summary, a sum-max MARL reward model with light-tailed components significantly out-performed computationally simpler models in reproducing empirical multi-syllable learning trajectories.

### Light-tailed sum-max intrinsic reward predicts the exclusion of a syllable from a song and its replacement with a call

The case of a bird failing to match a target (Figure 2G) raises an interesting question: what happens when there is a target that does not have any syllables in its reward range, *and* a syllable that is not within the reward range of any target? In such a case, our model predicts that the syllable would not shift from its mean (and in effect be dropped from the learning process), and that a nearby call could shift to the unoccupied target (since there are no syllables in that target’s reward range). We re-inspected the data of the bird shown in Figure 2G to search for a call performed in the acoustic vicinity of the unmatched target but found that no such call was present. Nevertheless, the possibility of a song syllable being left out in favor of a call triggered our interest and we decided to further investigate this prediction of our model.

We presented juvenile male zebra finches (n = 6) with a learning task in which one syllable in the target song is shifted up or down by 4 semitones with respect to a syllable in the bird’s current song (*AB* → *AB*^±4^; Figure 3A). In this experimental situation, there is a single off-target syllable in the bird’s song, *B*, and a single unmatched target in the tutor song, *B*^±4^, but the two are separated by a relatively large pitch difference. A hierarchical sum-max reward computation based on light-tailed components predicts that song syllable *B* is outside the reward range of the target syllable *B*^±4^ (Figure S5A) and therefore, that it will not shift to match the target. Consequently, if the bird’s repertoire happens to contain a call *C* that is within the reward range of the target, the call will shift to match it instead. In contrast, normal and heavy-tailed versions of the model predict that song syllable *B* is within the reward range of a 4-semitone-shifted target, and therefore will shift to match it, even if there is an acoustically closer call in the bird’s repertoire (Figure 3B), since syllables are prioritized over calls.

**Figure 3:**
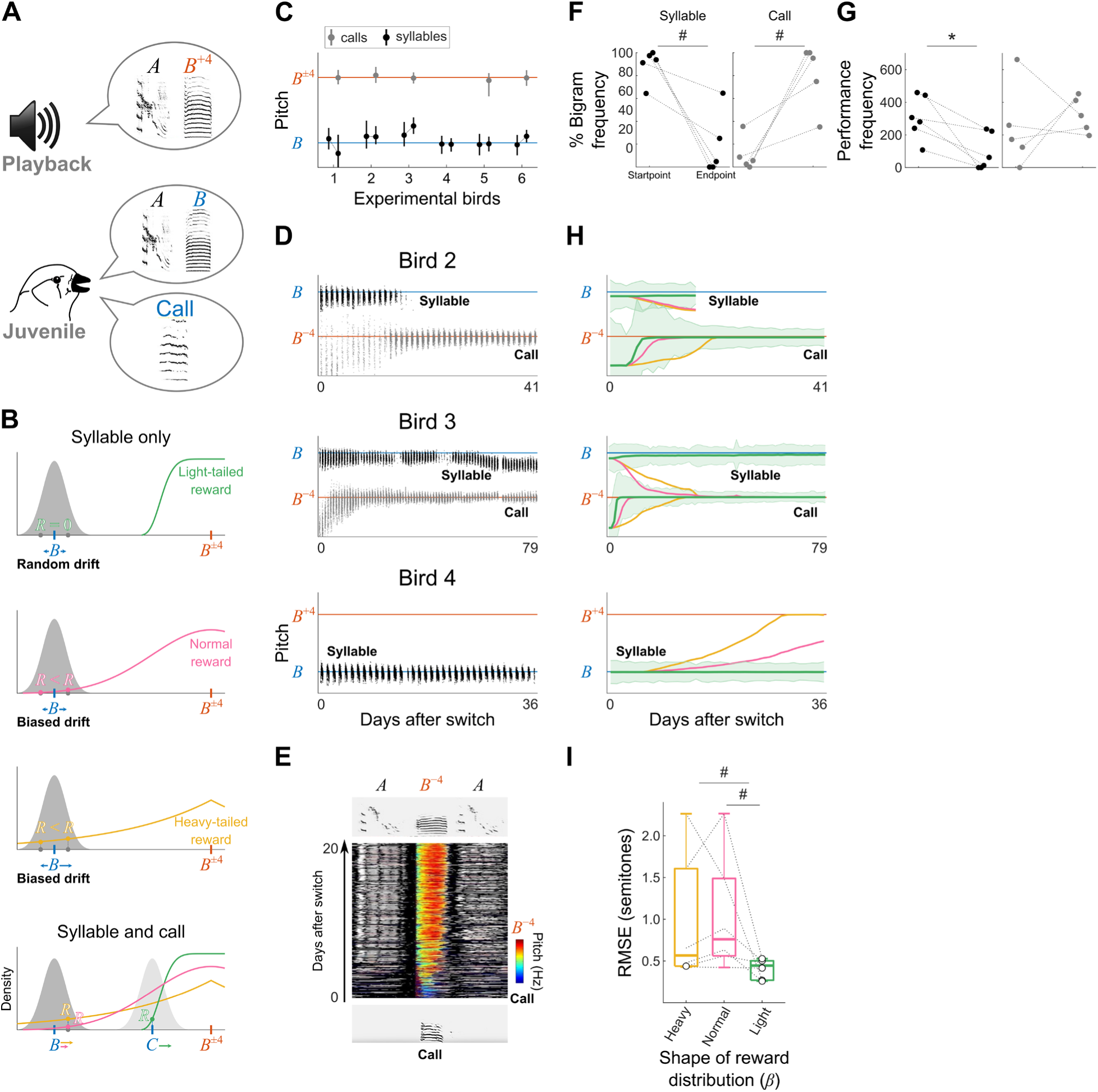
Light-tailed sum-max intrinsic reward predicts the exclusion of a syllable from a song and its replacement with a call. **A.** A syllable-matching experiment testing the prediction of a syllable being excluded from the song: a bird that has learned a two-syllable song *AB* is induced to learn a new song *AB*^±4^ where the target *B*^±4^ is 4 semitones above or below the song syllable *B* (an *AB* → *AB*^+4^experiment is shown). The bird’s repertoire includes song syllables and calls. **B.** Predictions of light-tailed, normal, and heavy-tailed versions of a MARL sum-max reward model for the experiment in **A**. Performance distribution of song syllable *B* (grey) and reward distributions centered on the target *B*^±4^ (colors as in Figure 1D) are shown. In a model based on light-tailed reward components, syllable *B* will not shift towards the target even if it is the most acoustically similar element to *B*^±4^ (top), because it is outside of the reward range. If a call in the bird’s repertoire is acoustically closer to *B*^±4^ (bottom, light grey), that call will shift to match the target. In contrast, normal and heavy-tailed versions of the model predict that syllable *B* will shift to match *B*^±4^ in any case, even if there is an acoustically closer call (middle and bottom). **C.** Results across experimental birds (n = 6) trained with the task in **A**. Notation as in Figure 2D. **D.** Learning trajectories of three experimental birds; from top to bottom: Bird 2 matched target *B*^−4^ with a call and stopped performing syllable *B*; Bird 3 matched target *B*^−4^ with a call and did not shift the pitch of syllable B; Bird 4 did not shift the pitch of syllable B, and did not recruit a call to match target *B*^+4^ (see also Figure S5B). **E.** Stack plot of consecutive instances of a call that shifted to match the target *B*^−4^ in bird 2 (**D**). Colors, pitch in call instances; greyscale, Wiener entropy in neighboring syllables. Sonograms show examples of the call performance at experiment start (bottom) and end (top). The call, initially performed outside the song motif (bottom), was incorporated into the song (top). **F.** Left, performance frequency of bigrams containing song syllable *B* and other motif syllables (% of total bigrams) at experimental startpoint and endpoint across birds. Right, same as left for the call that shifted to match target *B*^±4^ in 5 of the 6 experimental birds. Syllable-containing bigrams decreased from start to endpoint of learning, while call-containing bigrams increased, indicating that the call was incorporated into the song motif while syllable *B* was dropped out (p = 0.0625, Wilcoxon signed-rank test). **G.** Performance frequencies of syllable *B* (left) and the call (right) that shifted to match target *B*^±4^ (3-day average of total syllables performed) at the start and endpoint across birds. Syllable performance decreased significantly (p = 0.0312, Wilcoxon signed-rank test) while call performance increased in 3 out of 5 birds. **H.** Bootstrapped pitch-trajectory distributions of the syllable (top) and call (bottom; 2 out of 3 birds) of the three birds shown in **C**. Colors as in Figure 2E. The shaded area represents the median ± 50% CI (see also Figure S5C). Note that under normal and heavy-tailed models, the syllable and the call can converge to the same target, which is possible when two actors are within the reward range of a single and common target (also see Discussion). **I.** Root-mean-square error (RMSE) between experimental and bootstrapped syllable learning trajectories. Open circles represent the RMSE of the best reward model per bird. Light-tailed reward shows a tendency to improve goodness-of-fit compared to heavy-tailed and normal rewards (#: 0.05 < p < 0.1, Benjamini-Hochberg corrected Wilcoxon signed-rank test).

None of the six experimental birds shifted song syllable *B* towards target *B*^±4^ (Figures 3C-D and S5B). In 5 out of 6 birds, *B*^±4^ was instead matched by a call (or another sound originally performed outside of the song motif), which was initially acoustically closer to the target (e.g., Figure 3D, birds 2 and 3). In all 5 cases, the call was fully or partially incorporated into the song motif (Figure 3E), replacing song syllable *B*. This was evident from an increase in the frequency of transitions between the call and motif syllables, and a concurrent decrease in the frequency of transitions between song syllable B and other motif syllables (Figure 3F; cf. Figure S2E). In addition, the performance rate of syllable B decreased (Figure 3G), with two of the birds stopping performing syllable *B* altogether (bird 2, Figure 3D, and bird 5, Figure S5B). We conclude that syllable *B* was functionally dropped from the song motif, and its function was transferred to a call. In the single case where no call shifted to match target *B*^±4^, the bird continued performing syllable *B* at the original pitch (bird 4, Figure 3D).

We tested how well a light-tailed sum-max model fits empirical learning trajectories, compared to normal and heavy-tailed models, by simulating learning trajectories of both syllables and calls (Figures 3H-I and S5C-D). All simulations used previously estimated parameters (Figure 1), without further adjustments to the new data except for the performance distributions’ initial mean pitches and daily variances which we estimated in each bird separately. The light-tailed model was superior to normal and heavy-tailed model variants, returning smaller errors between simulated and empirical learning trajectories in 5 out of 6 birds (Figure 3I), and replicating the failure of the bird that did not have a call at the acoustic vicinity of the *B*^±4^ target to accomplish the learning task (bird 4, Figure 3H).

In summary, the experimental outcomes support the predictions of a light-tailed sum-max reward computation with prioritization of syllables over calls. These results demonstrate a surprising feature of vocal learning in zebra finches, which is a direct consequence of a competitive and light-tailed reward computation: an intrinsic reward is not necessarily contingent on all syllables a bird is performing, but only on those syllables that are sufficiently similar to at least one syllable in the target song. Syllables that are too dissimilar to contribute to the reward computation end up being dropped (fully or partially) from the song motif. These findings underscore a considerable modularity and flexibility of zebra finches’ developing songs and raise the interesting question of the adaptive value of such modular vocal development in birds’ complex natural social environment.

### Predicted tuning of error-encoding dopaminergic VTA neurons during the learning of a syllable inventory

The intrinsic inventory reward in our model is fed into a computation of the RPE *δ*, which tracks deviations from expected reward (Figure 1B and Materials and Methods). Assuming that *δ* is proportional to the putative firing rate of dopaminergic neurons projecting into the avian song system, our model provides predictions for dopaminergic firing in juvenile birds learning a multi-syllable inventory (Figure 4). We present our model’s predictions in a juvenile bird at the onset of learning in each of our three experimental scenarios: Learning a single target syllable shifted by 2 semitones from a syllable in the bird’s song (Figure 1D), leaning two different 2-semitone-shifted targets (Figure 2C), and learning a 4-semitone-shifted target (Figure 3A).

**Figure 4:**
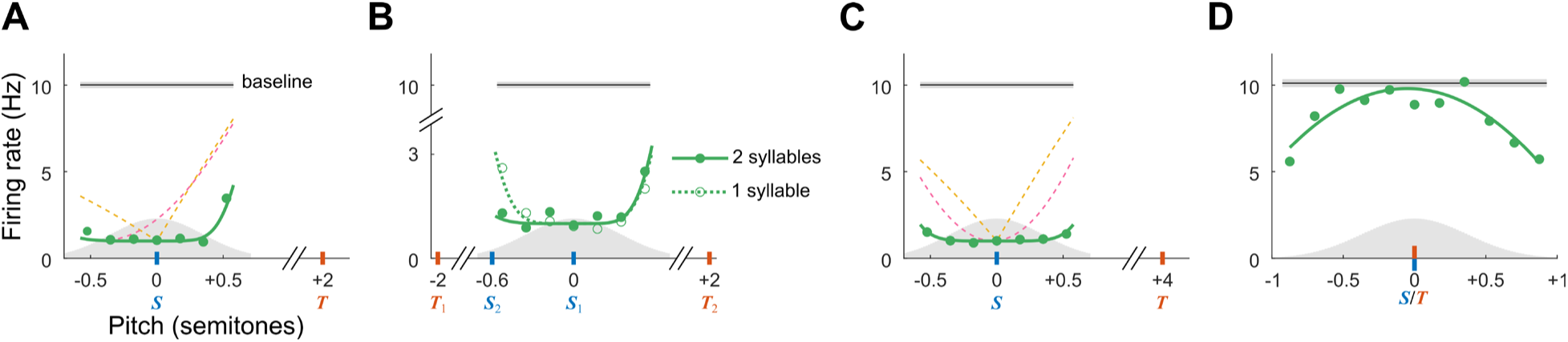
Predicted tuning of error-encoding dopaminergic VTA neurons during the learning of a syllable inventory. **A**. Predicted VTA tuning as a function of pitch in the case of a target (*T*, orange) shifted by 2 semitones with respect to a song syllable (*S*, blue). Green, theoretical (solid line) and simulated (filled circles) VTA tuning predicted by a light-tailed reward. Dashed lines, theoretical VTA tuning predicted by the heavy-tailed reward (yellow) and normal reward (magenta). **B.** Predicted VTA tuning in the case of two targets (*T*_1_ and *T*_2_). When a single syllable *S*_1_ is at a distance of 2 semitones from both targets, VTA tuning is symmetrical (dotted line, empty circles). The presence of a second syllable *S*_2_ between *S*_1_ and *T*_1_ breaks this symmetry, resulting in the VTA neuron firing more following instances of *S*_1_in the vicinity of *T*_2_, than instances in the vicinity of *T*_1_. **C.** Same as A for a 4-semitone-shifted target. VTA-neuron simulations in **A-C** are based on 2000 pitch values drawn from a normal performance distribution (grey shading) with a typical standard deviation at the onset of learning (*σ_B_* = 0.35 semitones). Firing rates, averaged in bins of size 0.5*σ_B_*, are in the range 1 to 10 Hz, where the higher end corresponds to the on-target, baseline firing rate. The hypothetical neuron emits Poisson spikes at a rate that scales linearly with RPE within a 150 msec window, beginning 50 msec after syllable onset. Baseline shows mean ± SEM. **D.** Predicted VTA tuning when a syllable matches a target, but the mean syllable pitch continues to drift slowly around the target due to circadian variations. The simulated VTA tuning (circles) predicted by a light-tailed reward is well fitted by a parabola (green line). Absence of pitch drift would result in a flat tuning (gray line). Simulation details as in **A**-**C**.

In the case of learning a single 2-semitone-shifted target (Figure 4A), the light-tailed reward model predicts selective tuning of VTA neurons to pitch values at the target-adjacent tail of the performance distribution (i.e., target selectivity) so that syllable pitch is driven in the direction of higher firing rate, i.e., smaller deviation from the target. However, target selectivity is restricted to a small range of pitch values at the tail, where the performance distribution overlaps with the light-tailed reward distribution. This means that VTA tuning is mostly insensitive to variations in pitch difference Δ between syllable and target pitch. This is consistent with reports of weak sensitivity of VTA response to the magnitude of negative RPEs upon external reinforcement^8,23,24^, which comes here as a direct consequence of the intrinsic light-tailed reward. The heavy-tailed and normal reward models also predict target selectivity of VTA neuron responses but fail to account for insensitivity to variations in Δ.

In the case of learning two different 2-semitone-shifted targets (Figure 4B), VTA tuning is expected to depend not only on the performance distribution of the currently sung syllable, but also on the distributions of other syllables (or calls) in a bird’s repertoire: The absence of competing syllables (i.e., of syllables that are within the reward range of one of the targets) would result in a symmetric VTA tuning (although in practice, a syllable would always be slightly closer to one of the targets, resulting in a slight asymmetry). In contrast, a competing syllable proximal to one target would break this symmetry, rendering the VTA neuron more selective to the other target.

In the case of a 4-semitone target (Figure 4C), the light-tailed reward model (in contrast to the heavy-tailed and normal reward models) predicts lack of target selectivity due to the reward being practically zero. This would lead to the syllable pitch drifting randomly around its mean (Figures 3B-D). Note that the zero-reward implies that the RPE *δ* ≈ −*Q*, which means the firing rate increases slightly with increased distance from the mean.

Finally, when a syllable matches a target (Figure 4D), i.e., in adult birds that have learned their song, the model predicts a flat pitch tuning curve due to absence of RPE. In reality, however, small pitch fluctuations^25^ are predicted to lead to the emergence of inverted-U tuning because the intrinsic reward decreases away from the target.

In summary, our model predicts that the syllable-related (or call-related) tuning of VTA neurons depends dynamically on the pitch difference between the vocalization and the target it is closest to. This leads to both asymmetric (Figures 4A-B) and symmetric (Figures 4B-D) tuning curves in dopaminergic neurons, and the symmetric curves can be either concave (Figure 4C) or convex (Figure 4D). Inverted-U and asymmetric tuning curves as in Figures 4A and 4D have been reported in adult birds^20^, but our predictions on tuning in juveniles including the dependence on multiple proximal syllables and calls (Figure 4B) remain to be explored.

## Discussion

Our work shows that an inventory of actions underlying a combinatorial behavior — birdsong — can be learned by a set of independent RL actors in a self-reinforced manner from an intrinsic scalar reward. By fitting alternative RL models to empirical developmental trajectories, we were able to infer and test the properties of this intrinsic reward, rejecting sequential-order-dependent models in favor of a greedy and context-independent model. This latter model assumes independence on three separate levels: among actors in the action system, among critics in the value system (see Materials and Methods), and among reward components in the reward system.

We believe that the independence and modularity among actors, critics, and partial rewards promotes the expansiveness and flexibility of inventory learning, making it easily scalable. Adding a target to the inventory turns into the simple problem of computing its associated reward and summing it up with the other partial rewards. The same goes for the removal of a target, which translates into elimination of that partial reward from the computation. The flexible goals this architecture supports may be an evolutionary adaptation for coping with changing targets in a dynamic environment and could explain the ability of juvenile zebra finches to combine syllables from multiple tutors under natural breeding conditions^26^.

Computationally, however, a greedy and context-independent sum-max reward seems dauntingly complex; especially in comparison to previous actor-critic models^2,3^, which assumed that syllables are learned within the sequential context of the song motif rather than independently of it (but see Troyer and Doupe, 2000^27^). In principle, a sum-max reward circuit (Figure 2B) needs to perform all possible pairwise comparisons at each time step – a total of *N* × *M* maximum operations per song motif, each over a set of *N* terms (for *M* targets and *N* syllables/calls). Although this seems like a daunting number for large inventories, the computation is quite manageable in practice, since among these *N*^2^ × *M* terms only at most 2*N* × *M* are distinct — the *N* × *M* terms comparing the targets with currently sung syllable instances and the *N* × *M* terms comparing the targets with the means *S_i_* (and *C_i_*) of the motor programs for syllables and calls not currently sung. Furthermore, since the pairwise comparisons are subserved by light-tailed reward distributions, the seemingly many terms in the inventory reward computation may further boil down to only a few nonzero ones associated with proximal syllable-target pairs. Thus, light-tailed reward distributions may promote efficiency.

In terms of neural architecture, a sum-max reward circuit can be implemented by a matrix-like network of *N* × *M* pairwise comparator neurons (each neuron corresponding to one pairwise reward component, Figure 2B). These neurons would have light tailed tuning curves, meaning that they would be tuned to small mismatches between a syllable and a target, but would be unresponsive to large mismatches. Biophysically, each pairwise comparator neuron would represent information about a target syllable *T_i_* and the mean of a motor program *S_j_* (or *C_j_*) either in terms of its neural activity, the synaptic inputs it receives, or the strength of its synapses. In addition, comparators would receive auditory input *P_j_* corresponding to feedback from a syllable instance just sung, which would replace the representation of the syllable’s mean performance *S_j_*. To implement the sum-max computation, the comparator neurons could be arranged in parallel winner-take-all modules, each of which computes a max operation with respect to a target via inhibitory synapses, followed by an output summation to obtain the inventory reward ***R*** (Figure 2B). All these computations are feasible with neural networks.

Where would this putative sum-max reward network, and the actors and critics that learn from it, reside in the songbird brain (Figure 5)? The comparator network computing the inventory reward *R* and the critics which compute the expected reward *Q* for each actor must reside upstream of VTA where the difference between these two signals (the RPE *δ*) is computed^8^. Candidate sites for the inventory reward network are the ventral pallidum (VP) and its afferents, given that VP input to VTA is of positive valence^28^ and therefore acts like the inventory reward *R* on *δ* (although another study found mixed signals in VP^29^). The critics would be located in the ventral intermediate arcopallium (AIV) and its afferents, given that AIV input to VTA^30^ is of negative valence^28^ and so acts like *Q* on *δ*. Both the VP and the AIV should in such a case receive auditory feedback from the syllable or call instance just sung *P_j_*, since this input is necessary for the computation of both *R* and *Q*. This assumption would agree with observations that neurons in VP and AIV are highly sensitive to distorted auditory feedback during singing^31^.

**Figure 5:**
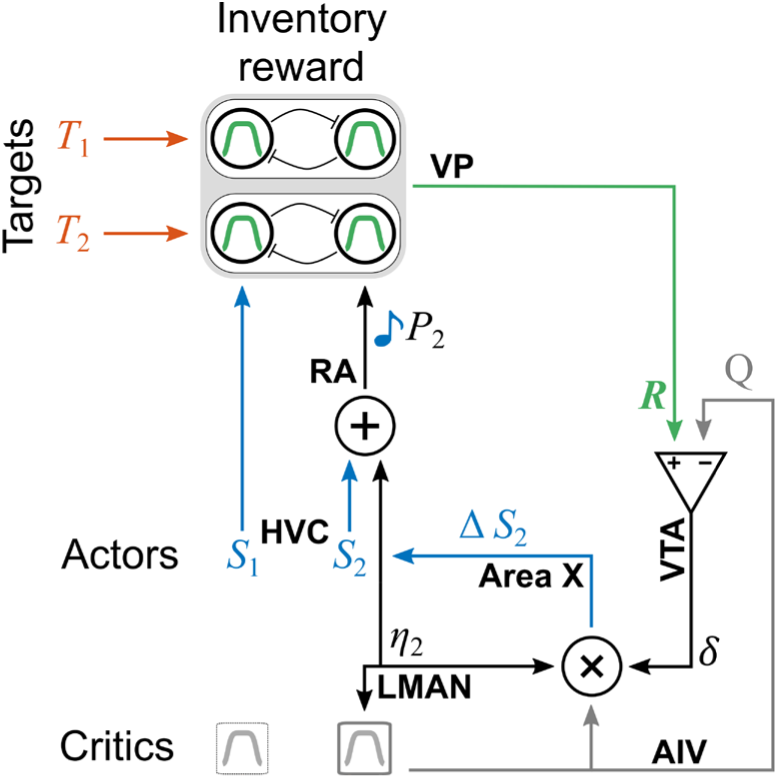
Potential mapping of the MARL model for inventory learning onto the songbird brain. Signals in our model (*S*, *P*, *η*, ***R***, *Q*, *δ*) are depicted as arrows and are labelled by the hypothesized brain areas (bold) where such signals can be found (implying the signals are computed in that area or upstream).

Since the premotor nucleus HVC controls the learned components of song, the lateral magnocellular nucleus of the anterior nidopallium (LMAN) is involved in vocal exploration, and both project to the robust nucleus of the arcopallium (RA), we suggest that the actors are distributed across HVC, LMAN, and RA. HVC would generate *S*, i.e., stable motor memories of each syllable or call^32^. The individual syllable/call instances *P* performed by RA would be the sum of two components: *S* (or *C* for calls) plus a more variable component *η* from LMAN^33^. This latter signal would be relayed as an efference copy to the dopaminergic recipient Area X where the learning itself could take place, in agreement both with a previous hypothesis^33^ and corroborating anatomical evidence^34^. The update Δ*S* computed in Area X would be consolidated in RA and HVC in a process that we do not explicitly model^35^.

The experimental paradigm and the model presented here contain some simplifications that could be broadened in future studies: First, we studied pitch – mainly because of our ability to selectively manipulate this acoustic feature. It remains to be seen how this approach generalizes to other sound features. Presumably, the pairwise reward components, as well as the critics, would need to be tuned to acoustic features other than pitch. Secondly, we made the simplifying assumption that the reward components and the critics have an identically shaped tuning to pitch differences. This convenient simplification may be relaxed in a more powerful version of the model, in which critics acquire their tuning curves via function approximation methods. Thirdly, we currently treat syllables as the basic, or smallest, units of learning, but in fact zebra finches can selectively adjust individual sub-syllabic elements towards increased target match^22^. This calls for tutoring experiments targeting the process of learning sub-syllabic notes, and a corresponding update of an intrinsic reward model to include the evaluation of sub-syllabic structure. Fourth, despite the maximum operation we used, the greediness our model implements is not strict (as in a soft winner-take-all) and two actors can in principle converge to the same target when no other target is around (for example, Figure 3G, middle). This possibility in our model arises from the stepwise updates Δ*S* of performance means, when two roughly equidistant means are pulled to the same target in an alternating manner. Although a light-tailed model makes this scenario rare by limiting the pitch range in which it can occur, such violation of greediness remains to be tested in future experiments.

Our findings suggest that the evolutionary driving force that has shaped birds’ learning strategy may have been efficiency – to be able to reach a set of targets while minimizing both the total amount of vocal change and the computational load. Given its efficiency and flexibility, the learning algorithm we discovered in songbirds may have evolved in other learned combinatorial behaviors, including the learning of speech sound inventories of human languages.

## Author Contributions

Conceptualization, H.T., A.T.Z., O.T., R.H.R.H., and D.L.; Methodology, H.T., A.T.Z., R.H.R.H., and D.L.; Software, H.T.; Formal Analysis, H.T.; Investigation, H.T., and D.L.; Resources, O.T., R.H.R.H., and D.L.; Data Curation, H.T., and D.L.; Writing – Original Draft, H.T., R.H.R.H., and D.L.; Writing – Review & Editing, H.T., A.T.Z., O.T., R.H.R.H., and D.L.; Visualization, H.T., R.H.R.H., and D.L.; Supervision, R.H.R.H.; Project Administration, R.H.R.H., and D.L.; Funding Acquisition, H.T., O.T., R.H.R.H., and D.L.

## Materials and Methods

### 1. Experimental design

Animal care and experimental procedures were conducted in accordance with the guidelines of the US National Institutes of Health and have been reviewed and approved by the Institutional Animal Care and Use Committee of Hunter College.

Part of the experimental data presented here (all 2-semitone learning experiments except for two new birds) was previously published^16^ and used here to infer and test model parameters. Male zebra finches were bred at Hunter College and reared in the absence of adult males between days 7–30 post hatch. Afterwards, birds were kept singly in sound attenuation chambers, and continuously recorded. From day 33–39 until day 43, birds were passively exposed to 20 playbacks per day of a tutor song (source), occurring at random with 0.005 probability per second. On day 43, each bird was trained to press a key to hear song playbacks, with a daily quota of 20. Once birds learned the source song, we switched to playbacks of a different tutor song (target). Learning of the source was assessed by quantifying the percent similarity (Sound Analysis Pro^36^) between the bird’s song motifs and the source model motif in 10 randomly chosen song bouts per day. We considered the source song as being learned when the similarity to the model was at least 70%. Since the sensitive period for song learning in zebra finches ends around day 90–100 post hatch, we included only birds that learned the source before day 71 in the experiments. Recording and training were done using Sound Analysis Pro^36^ and continued until birds reached adulthood (day 99–158 post hatch). At these ages, males are sexually mature, and perform a crystalized song motif that remains unchanged for the remainder of their lives^37^.

Source and target song models were synthetically composed of natural syllables. Harmonic syllables in the source songs were pitch-shifted by 2 or 4 semitones in the target songs using GOLDWAVE (v. 5.68, www.goldwave.com). Each playback of a model included two motif renditions. To control for model-specific effects, we varied baseline pitch, and pitch shift direction across experimental birds.

### 2. Data analysis

We performed song feature calculation and clustering of syllables and calls using Sound Analysis Pro^36^ (see Lipkind et. al., 2017^16^). All other analyses were performed in MATLAB (Mathworks Inc). Unless mentioned otherwise, significance levels were adjusted using the Benjamini-Hochberg procedure^38^ to correct for multiple comparisons.

*Calculating median pitch of syllable instances*: We used the following procedure to prepare pitch traces for statistical analysis and model fitting. Following syllable clustering, median pitch values of syllable instances *P* were converted from hertz to semitones using the formula:

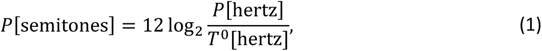

where *T*^0^is the source pitch (i.e., the pitch of the syllable in the source song that we manipulated in the target song). The semitone scale allows us to express our results in a common reference frame in which the source pitch is zero. In order to minimize biasing effects of clustering errors on model fits, we excluded renditions with pitch more than 3 median absolute deviations from the daily median pitch (leading to elimination of less than 0.2% of data points).

*Detrending daily pitch distributions*: Some analyses required day-wise detrending of pitch values to factor out the effects of learning and circadian patterns. We performed day-wise detrending by subtracting a 30-rendition pitch running average within each day from that day’s pitch values (analysis was robust to window sizes of 50, 75, and 100 instances). This detrending procedure was used for calculating daily and sub-daily pitch standard deviations (see Figure S2F).

*Characterizing daily pitch distributions*: We first restricted the distribution analysis to stable days: we assessed pitch stability across consecutive day pairs (i.e., days with no learning or no random drift) by comparing medians of pitch distributions (uncorrected Mann-Whitney U test; *α* = 0.001). On the set of stable day pairs, we found pitch distribution to be nearly always Gaussian: The daily distributions (with more than 10 instances) of stable pitch were normally distributed on 730/778 days = 94% (uncorrected Kolmogorov-Smirnov test on standardized distributions; *p* > 0.05). The same was true when we tested on all days, not just the stable ones, after day-wise detrending (uncorrected Kolmogorov-Smirnov test on standardized distributions *p* > 0.05 in 1214/1276 days = 95%). This justifies our choice of the normal performance distribution for the actors in our RL model (see below).

### 3. Reinforcement learning model

Our RL model consists of three components: actors that are responsible for generating vocalization (syllable and call) instances, an intrinsic reward ***R***, and a critic that estimate the expected reward *Q* of an instance. We first describe the model of a single actor-critic pair suitable for learning a single target syllable.

#### 3.1 Single-actor model

At time *t*, the actor generates a syllable/call instance with pitch *P*(*t*) = *S*(*t*) + *η*(*t*), where *η* is drawn from a normal distribution,

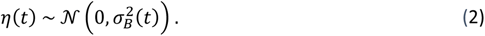

*S*(*t*) is the mean performed pitch and *σ_B_*(*t*) is the (time-dependent) standard deviation that sets the extent of behavioral variability. The syllable instance is followed by an intrinsically generated reward signal *R*(*t*) ≔ *R*(*P*(*t*), *T*) that depends on both the produced pitch *P*(*t*) and the target pitch *T*. We assume that *R*(*t*) is symmetric around *T* and is a decreasing function of the absolute pitch difference Δ(*t*) ≔ |*P*(*t*) − *T*|. Because the functional form of *R*(*t*) is *a priori* unknown, we introduce a family of hypothetical intrinsic reward functions, i.e., we model reward as a generalized normal distribution^39^,

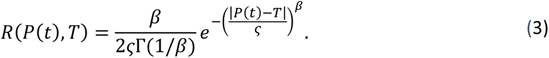

The parameter *β* sets the shape of the reward distribution around the target: larger values of *β* correspond to distributions with lighter tails with the reward density concentrating more around the mean. Special cases of this family of distributions are normal and Laplace distributions given by *β* = 2 and *β* = 1, respectively. The positive scale parameter *ς* controls the width of the reward distribution. The reward standard deviation *σ_R_* depends on both *β* and *ς*, 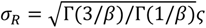.

Learning is driven by the bird’s attempt to maximize reward via minimizing the square discrepancy *δ*^2^(*t*) between actual reward *R*(*t*) and expected reward *E*(*R*(*t*)|*P*(*t*)),

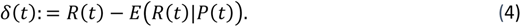

The variable *δ*(*t*) is also referred to as the reward prediction error (RPE), which is commonly assumed to be related to the firing rate of dopaminergic neurons. To estimate the expected reward *E*(*R*(*t*)|*P*(*t*)) associated with a vocalization *P*(*t*), we introduce a parametric function *Q*(*t*) ≔ *E*(*R*(*t*)|*P*(*t*)), referred to as the critic. We give the critic the same functional form as the reward distribution *R*(*t*), but we center its peak on the average pitch *S*(*t*) instead of the target *T*,

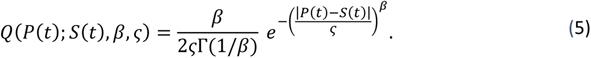

Our assumption that *R* and *Q* have the same functional form is plausible because presumably both *R* and *Q* are computed by neural circuits that have co-evolved to support song learning.

We derive the learning rule by minimizing the mean-square error 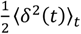 using stochastic gradient descent, which is the conventional approach to continuous-action RL^40^:

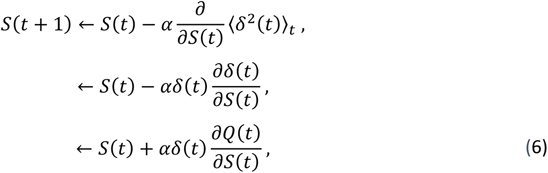

where *α* > 0 is the learning rate.

Inserting Eq. 5 into Eq. 6 leads to the following Rescorla-Wagner-type iterative update rule for the mean pitch *S*(*t*):

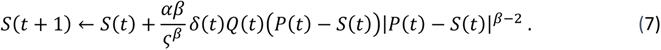

According to this rule, updates of *S*(*t*) mostly point in the opposite direction of *P*(*t*) (*δ* is rarely positive) and are proportional in magnitude to the RPE *δ*(*t*), which biases the drift of *S* towards *T*. During learning, as the RPE converges toward zero, *Q* approaches *R*, and the syllable’s mean pitch *S*(*t*) approaches the target pitch *T*.

It is important to note that although, by construction, the goal of our model is to minimize the square RPE (*δ* model), an RPE is not needed in principle. Instead, we could have chosen the goal of directly maximizing the intrinsic reward (as in policy-gradient learning^41,42^) Such a model would be simpler in fact, as it would not require a critic for computing an expected reward, which would free up resources. Under an *R* model, assuming that dopaminergic responses in VTA encode *R* instead of *δ*, the experimental predictions would be high firing rates in VTA neurons of adult birds with an accomplished song (i.e., triggering maximal reward) and light-tailed singing-related pitch tuning curves (i.e., response tuning curves with the same shape as a light-tailed reward component).

However, the rather low average song-related firing rate (13 Hz) of Area X-projecting VTA (VTA_X_) neurons observed in adults^43^ does not strongly support this view, suggesting instead that dopaminergic neurons encode a difference signal, as predicted by a *δ* model. Neither is the tendency of VTA neurons to display inverted-U tuning curves centered on a single auditory feature^20^ in unequivocally strong support for the R-Learning model; in fact, we found a similar behavior under the *δ* model when we let the mean pitch of an actor slowly drifts around its target, which also led to bell-shaped tuning (Figure 4D). In summary, it is currently not possible to unequivocally arbitrate between the *δ* and R-Learning models, the R-Learning model may be simpler but the *δ* model may map better onto known firing properties.

#### 3.2 Multiple-actor model

Next, we consider the case of many actors that learn to produce a multi-syllable inventory. We introduce an independent actor for each of the juvenile’s vocalization types. Actor *j* (*j* = 1, …, *N*) randomly produces vocalizations (a syllable or a call) drawn from a normal distribution with mean pitch *S_i_* and standard deviation *σ_B_*_,*i*_ (*t*). The *N* actors compete to fill *M* targets of pitch *T_i_* (*i* = 1, …, *M*). The actors learn from a common scalar *inventory reward R*, which agrees with RPE being signaled by a single neuromodulator, i.e., dopamine.

Our model must reflect the observation that birds make the minimal changes necessary to match the target inventory, i.e., vocalization assignments are greedy and independent of sequential context. To satisfy the latter constraint of independence, we write the inventory reward ***R*** as a sum (which is insensitive to permutations) of partial rewards,

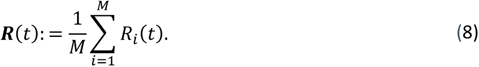

The partial reward *R*_*i*_ (*t*) associated with target *i* must reflect the maximum across individual actor-target pairwise comparisons. Naively, to implement a competition between actors over a target, we could choose a simple maximum operation across all actors^1^, since only one actor can score the partial reward associated with a given target, namely the one that produced the acoustically closest instance among all vocalization types in the inventory. However, to account for the prioritization of syllables over calls illustrated in Figures 1L and S4B, we define the partial reward *R*_*i*_ for target *i* as the **syllable-specific partial reward**,

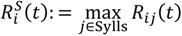

if the latter is positive (i.e., larger than some infinitesimal value *∈*), else we define it as the **call-specific partial reward**,

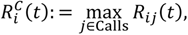

as follows:

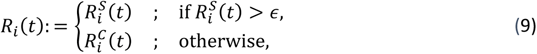

where ∈ = max(*R_ij_*) /1000 is a small number (see Figure S3A). According to this definition of partial reward, a distant syllable within the reward range of a target will win over a closer call, replicating the finding in Figures 1L and S4B.

We are left with needing to define *R_ij_*(*t*), the pairwise reward primitives and relate them to the estimated reward density *R*(*P*; *T*) of the single-actor model from the previous Section. The bottleneck to consider is that we can compute the maximum operations in Eq. (9) only after all arguments are known. To avoid the need of having to wait until all calls and syllables in the inventory have been produced, e.g., at the end of a song motif, we formulate a model in which rewards and error signals are computed upon production of every syllable and call instance with minimal requirement on short-term memory (see Section Motivation for Instantaneous Reward).

We assume that the reward primitive *R_ij_* (*t*) associated with the *i*’th target and the *j*’th actor depends on the vocalization instance *P_i_* (*t*) just performed and the syllable motor means *S_i_* (*t*) (*C_i_* (*t*) for calls) of non-performed vocalizations,

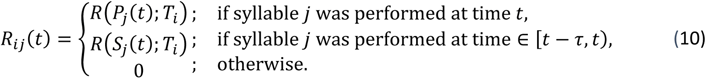

Only syllables and calls that were recently performed contribute to ***R***, which is encapsulated by the time window τ = 1 min, corresponding roughly to the average duration of a singing-and-calling period (see the distributions of call-song intervals in Figures S3C and S5E).

Note that context-independent inventory learning as we propose it here is also offered in the model of Troyer and Doupe^44^, where song performance is evaluated in parallel modules, one for each target syllable; however, these modules are combined via rich synaptic weights and so the overall evaluation in their model is far from being scalar in nature as the dopamine signal we model.

#### 3.3 Critics

For each actor *j*, we introduce a critic that tracks the expected reward *Q_j_* (*t*) ≔ *E*(*R*(*t*)|*P*_j_ (*t*)) associated with each syllable/call instance performed by that actor. The quality function *Q_j_*(*t*) = *Q*(*P_j_* (*t*); *S_j_* (*t*), *β*, *ς*) of the critic takes the same form as in Eq. 5. The parameters *β* and *ς* are common to all critics, but each critic has a distinct mean pitch parameter *S_j_*(*t*) (*C_j_*(*t*) for calls) that it inherits from the actor it is paired with. When actor *j* produces a vocalization, only critic *j* produces a non-zero expected reward. All other critics produce an output of zero: *Q_k_*_≠*j*_(*t*) = 0. It follows that the total expected reward *Q*(*t*) of a vocalization is simply the summed output of all critics, *Q*(*t*) = Σ*_k_ Q_k_*(*t*).

#### 3.4 Gradient descent learning rule

Given the intrinsic reward *R*(*t*) in Eq. 8 and the expected reward *Q*(*t*) as above, we define the RPE again as *δ*(*t*) = *R*(*t*) − *Q*(*t*) and minimize the mean square of that error, 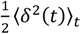, in a stepwise manner using stochastic gradient descent. We obtain an iterative update for the mean *S_j_* of an actor that is analogous to Eq. 7,

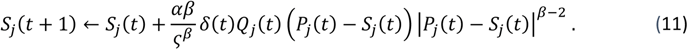

Thanks to the term *Q_j_*(*t*), we see that only the mean of the actor that generated a vocalization is updated, whereas all other actors are left unchanged.

Equation 11 is in essence a Hebbian-like learning rule that multiplicatively combines 3 terms: an RPE *δ* that is common to all actors and evaluates whether the performance is better or worse than expected, a critic *Q_j_* that restricts the update to the actor responsible for producing the vocalization, and an efference copy *η_j_* = *P_j_* − *S_j_* that provides information about whether the pitch of the current vocal instance was above or below its mean.

Such a triplet learning rule as in Eq. 11 has been predicted to exist in the basal ganglia of songbirds, and recent data provides anatomical support for its existence^45^. According to this line of work, there is convergence of three types of signals onto medium spiny neurons (MSN) in Area X of the songbird basal ganglia: an error signal (like *δ*) that stems from VTA, a timing signal (like *Q_j_*) from HVC that restricts learning to the right time in the song, and an efference copy signal (like *η_j_*) from the lateral magnocellular nucleus of the nidopallium (LMAN) that provides information about the current exploration (see Figure 5). These signals emerge naturally from our multi-actor model trained to minimize square RPE, providing anatomical support for our approach.

#### 3.5 Illustration of dynamic assignment

The competitive-cooperative reward computation in Eq. 8 and Eq. 9 entails *dynamic* assignments between vocalizations and targets, which we illustrate with a specific example. When multiple syllables are in the “reward range” of more than one target (Figure S3B, top), syllable-target assignments may initially vary from instance to instance, sometimes leading to an update of a motor program’s mean towards one target, and sometimes towards another. However, even small asymmetries in the reward magnitudes across different syllable-target pairings will lead to cooperative convergence of assignments that maximize inventory reward (Figure S3B, middle), and eventually to all targets being matched (Figure S3B, bottom). This dynamic process is reminiscent of a musical chairs game, in which pairings between players and chairs are not predetermined, but are resolved in a competitive-cooperative manner, eventually resulting in each chair being occupied by a single player.

### 4. Model Inference and Evaluation

We infer model parameters from experimental birds induced to shift the pitch of a syllable toward a target that differs by 2 semitones upwards or downwards (Figure 1; Eq. 7). We included experimental birds that were trained to shift the pitch of two different syllables towards two different targets separated by at least 6 semitones (birds shifted each of the source syllables to match a target without competitive interactions among the syllables). Overall, the data included 13 syllables from 8 birds. We excluded 3 out of 13 syllables that showed transient switching back to the source pitch after crossing the midpoint between source and target (see Figure S1A). Since were specifically interested in the part of a trajectory containing the learned shift toward the target, we extracted from each experimental dataset the learning subset of pitch values ***P_L_*** = {*P*(1), *P*(2), …, *P*(*t*), …, *P*(*L*)} from *D* days. These days contain the shift (identified from the day at which the pitch crossed the midpoint between source and target syllable), in addition to (at most) 3 consecutive days of stable pitch immediately before and after the shift (we worked with pitch data from fewer than 3 consecutive days either when the bird started shifting on day two post switch or when the experiment was stopped early such that stable pitch data was not available for 3 full days after the switch).

Model parameters are estimated from ***P_L_*** as follows. Juvenile birds in artificial tutoring experiments often show small deviations from a perfect imitation of the target song in addition to random daily fluctuations in pitch. Therefore, the mean pitch *S*(*t*) of a syllable (Eq. 7) does not necessarily begin at *T*^0^ on *t* = 1 and end at *T* on *t* = *L*, leading us to add the start and end pitch values *T*^0^′ and *T*′ as free model parameters. Data analysis also shows that pitch variance *σ_B_*(*t*) changes over days but is stable within a day (Figure S2F). We thus include an additional set of parameters ***σ****_B_* = (*σ_B_*_,1_, …, *σ_B,D_*), where *D* is the number of experimental days over which the dataset ***P_L_*** was collected, as described above. Birds display a variable latency between the switch to target song playback and the onset of pitch modification. In our model, this corresponds to a step-change in the mean of the reward distribution from *T*^0^′ (where *δ* = 0) toward *T*′ (where *δ* ≠ 0), initiating pitch modification. We therefore add an integer parameter *c* that defines the step-change index at which the pitch target in the model changes from *T*^0^′ to *T*′.

We infer all model parameters ***θ*** = {*α*, *β*, *c*, ***σ****_B_*, *σ_R_*, *T*^0^′, *T*′} including the shape *β* and standard deviation *σ*_*R*_ (or a subset of those parameters; see below) using maximum likelihood estimation:

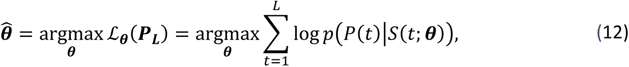

where 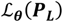 defines the log-likelihood that ***θ*** generates data ***P_L_***, and *S*(*t*; ***θ***) is the motor mean as a function of ***θ*** (see Eq. 7). The log-likelihood is maximized through nonlinear constrained optimization using MATLAB’s active set algorithm (fmincon). Parameters {*α*, *β*, ***σ****_B_*, *σ_R_*} are constrained within the interval [0, +∞), while *T*^0^′ and *T*′ are constrained to within 0.5 semitones from the true source and target pitch, *T*^0^ and *T*, respectively. The step-change index 1 ≤ *c* ≤ *L* is rounded to the closest integer after parameter inference. To avoid convergence to a local maximum, we start the maximization routine from different initial points along a grid in parameter space (except ***σ****_B_* which were initialized from day-wise detrended data; see above). Note that *σ_R_* is estimated from syllables only. Determining the standard deviation of the reward distribution of calls remains an outstanding problem.

We considered multiple model variants. We refer to the model containing the complete set of parameters ***θ*** as the full model. We also inferred the parameters of two constrained models where either the reward shape *β* alone or both *β* and the learning rate *α* were fixed (i.e., we infer ***θ*** = {*α*, *c*, ***σ****_B_*, *σ_R_*, *T*^0^′, *T*′} or ***θ*** = {*c*, ***σ****_B_*, *σ_R_*, *T*^0^′, *T*′}, respectively). In the latter case, we set *α* to 0.975, the median estimated value in the full model. When fixing *β*, we used a range of values corresponding to heavy-tailed (*β* = 1), normal (*β* = 2), and light-tailed (*β* = 4, 6, 8).

Assessing the goodness-of-fit using the Bayesian Information Criterion^46^ showed comparable results across different model variants (Figures S2A, S2C and 1H).

We further evaluated the models’ goodness-of-fit by simulating 500 pitch trajectories given the inferred model parameters and computing the root-mean-square-error (RMSE) between simulated and real data. This resulted in a distribution of RMSEs, D_RMSE_, per syllable that allows us to assess a model’s ability to generalize. To do so, we summarized the central tendencies of those distributions using Tukey’s trimean,

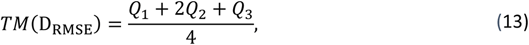

and compared model performance across different parameter settings over the entire set of syllables used for inferring parameters.

We found that the simplest model where both *α* and *β* were fixed significantly outperformed the more complex models (see Figure S2D), suggesting that BIC does not capture well the model’s ability to generalize. We therefore used the simplest model and its inferred reward standard deviation parameters (ten *σ_R_* values, corresponding to the 10 syllables in the training set in Figure 1) in all testing simulations (Figures 2 and 3, 2-semitone shift toward two potential targets and 4-semitone shift toward one target; 100 simulations per *σ_R_* value, amounting to 1000 simulations per bird in total). In these simulations, we specified other model parameters from the data directly, using the procedures described above. In case a syllable did not shift towards a target (e.g., Figures 2G and 3D), *c* was set to the time at which another vocalization in the bird’s repertoire (usually a call) started shifting toward the target. In the one case where no vocalization in the bird’s repertoire shifted toward the target (see Figure 3D, bird 4), *c* was set to its median value over the rest of the experimental dataset. After simulations, we used Tukey’s trimean described above, Eq. 13, to compare model performance for different reward shapes across each experimental dataset (Figures 1J, 2F and 3H).

### 5. Motivation for instantaneous reward

In a naive MARL version where all reward primitives are based on produced pitch, *R_ij_* = *R*(*P_i_*; *T_i_*), inventory reward can only be computed after all syllables and calls have been produced, e.g., at the end of a song motif. Such a model would thus require birds to maintain recent syllable and call instances in short-term memory. However, it is unclear whether the brain would choose such a cumulative strategy of computing reward rather than making an instantaneous estimate available after each syllable and call instance. A fine temporal signaling of inventory reward agrees with highly phasic dopaminergic responses, which can occur in the middle of a motif right after an aversive external stimulus or after the omission of a stimulus^8^. Also, dopaminergic activity seems to be roughly uniformly distributed during singing and there are no reports of its accumulation at motif endings, further suggesting a tight temporal correspondence between syllables and intrinsic rewards. Instantaneous reward signalling is also supported by studies involving experimental interference with birds’ auditory feedback in a manner contingent on syllable pitch, showing that interference must be closely time-locked to syllable performance to drive behavioral changes^47^.

### 6. Null models

As null hypotheses, we first adapted the model proposed by Fiete and colleagues^3^, where each actor *j* is assigned to the target *T_i_* in a sequential-order-dependent (SOD) manner (i.e., where vocalization *P_i_* (*t*) is temporally aligned to *T_i_* in the target song; see Figure S4E). In the SOD null model, reward at each vocalization instance is a function of the similarity between that instance and the temporally aligned target only, ***R*** = *R_ij_* = *R*(*P_i_*, *T_i_*). Reward in one instantiation (SOD 1) is binarized as in the original model^3^, where *R* = 1 when the absolute error (*P_i_* − *T_i_*) is below a similarity threshold. The similarity threshold is adaptive (using an exponential moving average of reward history) to assure that the actor is rewarded half the time. We give the critic the same functional form as the actor (Eq. 2), to assure that *Q_j_*(*t*) is centered at the mean *S_j_*(*t*) and is a differentiable function of the mean, enabling the derivation of a learning rule with stochastic gradient descent (as in Eq. 6). A second instantiation (SOD 2) uses the same light-tailed reward distributions as in the MARL model.

Another sum-max model (sum-over-actors model) consists of multiple actors as in the sum-max MARL model, but rather than actors competing over a target, targets compete over an actor by defining each partial reward associated with actor *j* as a maximum across pairwise comparisons between actor *j* and all the targets. This is followed by a sum over the partial rewards for all actors (see Figure S4C, right).

### 7. Alternative sum-max MARL model variants

Our findings based on the inventory reward model in Eq. 9 and Eq. 10 are robust to changes in model variant and model parameters. *A priori*, many models can signal inventory reward after every produced syllable. In another model variant we simulated, the partial reward *R_ij_*(*t*) associated with Target *T_i_* did not depend on the motor means *S_j_*(*t*) but instead it depended on the instance memories *P_j_*(*t*) of syllables and calls that were last performed within *τ*. Results were qualitatively unchanged under this instance-memory model, for both shorter and longer windows (from *τ* = 2 seconds to *τ* = 2 hours). Finally, note that instead of minimizing the square RPE in Eq. 6 and Eq. 11, it would also be possible to maximize reward via gradient descent (in which case no critic *Q* would be needed). Although such a model variant sounds appealing due to its simplicity and would be worthwhile further studying, we have chosen to simulate a variant based on minimization of RPE due to its hypothesized role underlying dopaminergic signaling.

## Supplementary Figures

**Figure S1:**
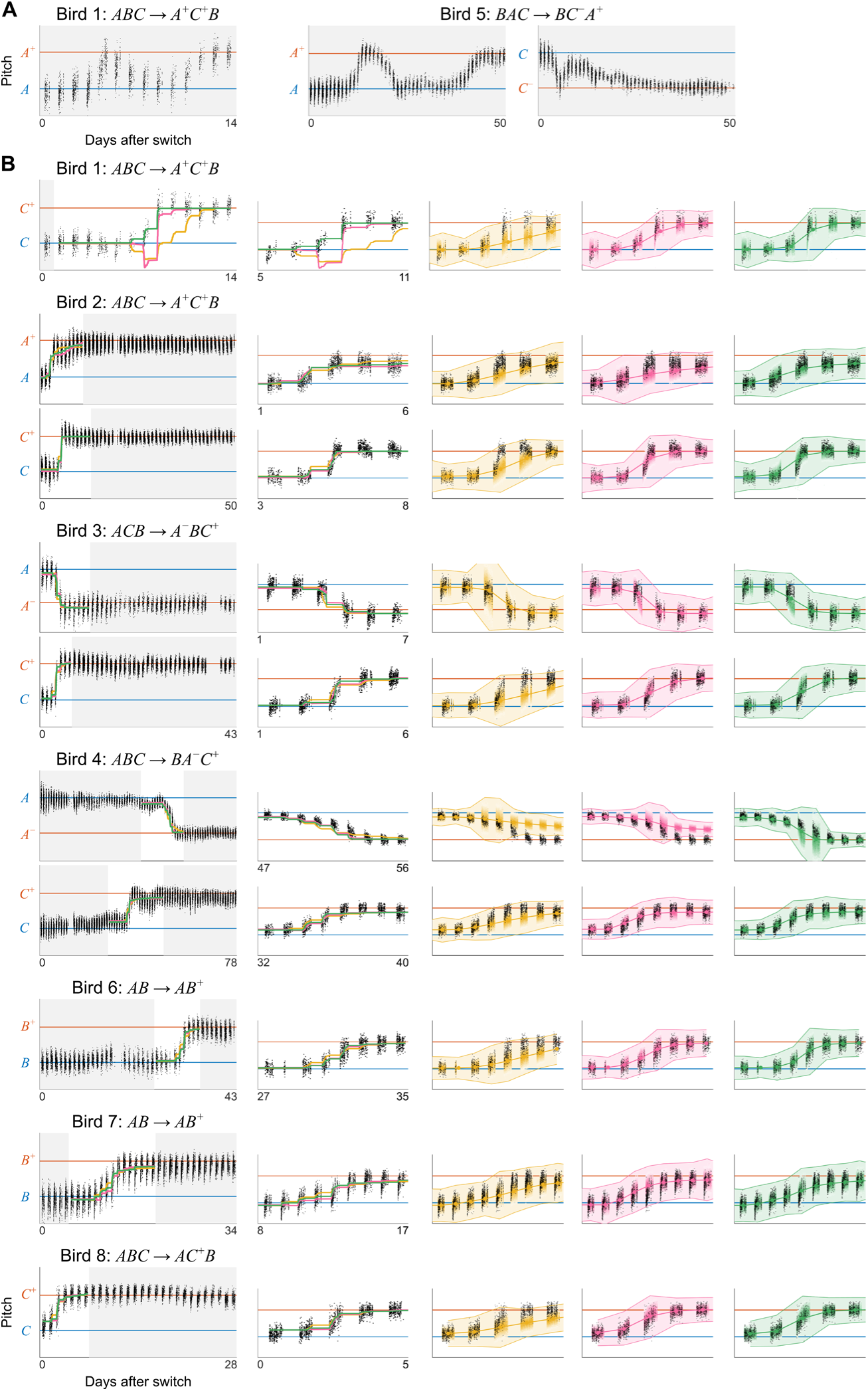
Observed, inferred, and bootstrapped pitch trajectories for all experimental birds tutored on imitation tasks requiring the matching of a target syllable shifted by 2 semitones. **A.** Syllable learning trajectories of birds trained with single-syllable learning tasks (Figure 1E, bottom), that were excluded from parameter estimation. Observed median pitch of consecutive syllable instances after the presentation of a new tutor song (day 0). **B.** Observed, inferred, and bootstrapped pitch trajectories for all syllables included in parameter estimation. From left to right: observed median pitch of consecutive renditions of the pitch-shifted syllables (black) after the presentation of the new tutor song (day 0), in addition to inferred mean pitch trajectories for different values of *β* (yellow, *β* < 2; magenta, *β* = 2; green, *β* > 2); zoom-in on the learning part of each pitch trajectory; Observed (black) and bootstrapped (color) pitch trajectories with shaded area representing median ± 50% CI of simulated trajectories for different values of *β*. Color gradient represents bootstrap density.

**Figure S2:**
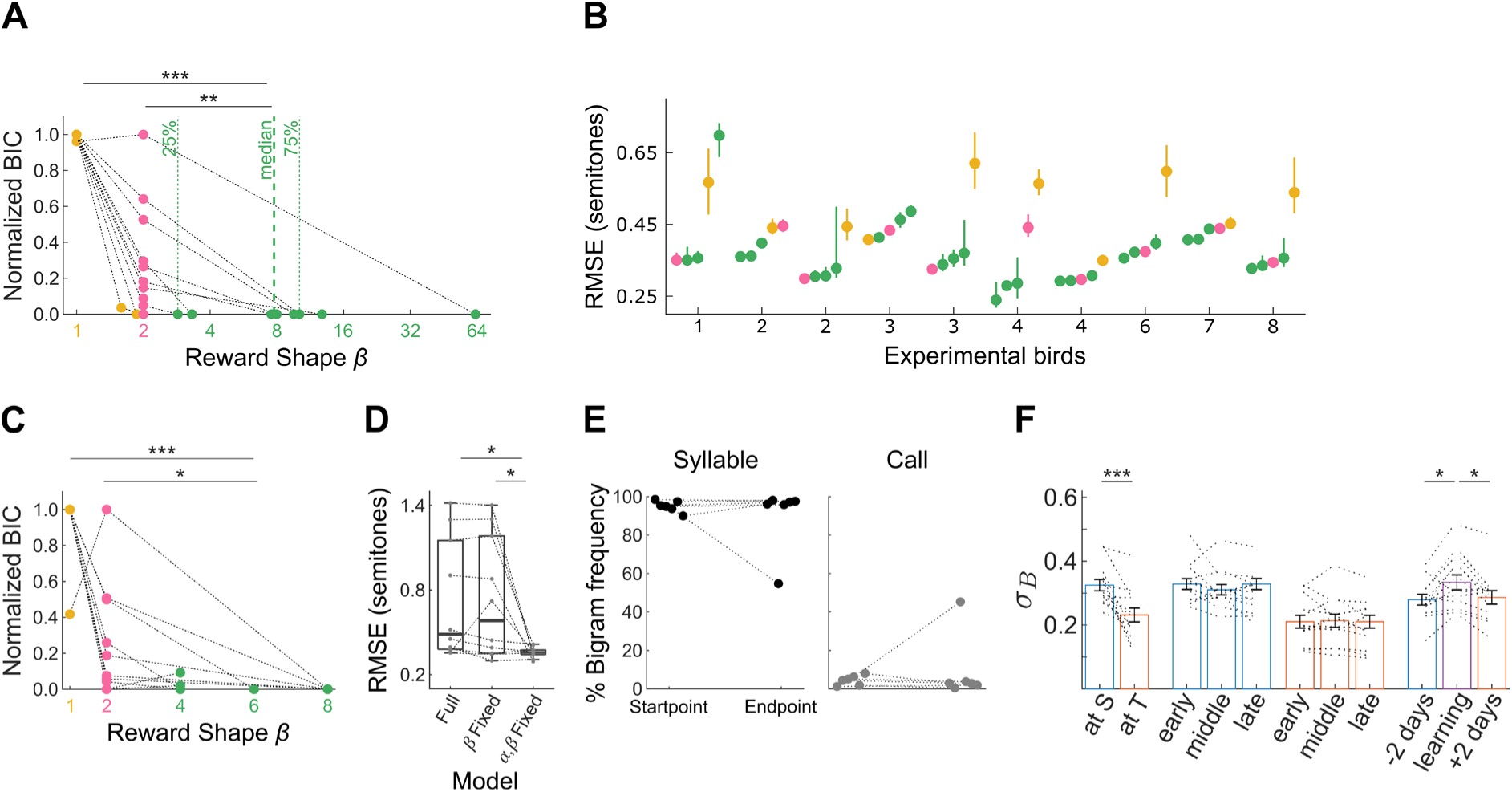
Model inference. **A.** Bayesian Information Criterion (BIC) for each fitted syllable (connected lines, n = 10) and shape (value of *β*). Shape is either fixed to 1 (heavy-tailed; yellow), 1 (normal; magenta), or inferred with maximum likelihood (full model, also learning rate *α* is inferred: 8/10 light-tailed *β* > 2; green; 2/8 heavy-tailed *β* < 2). BIC values for each syllable are normalized such that 1 corresponds to the worst and 0 to the best fit. Light-tailed reward improves data likelihood compared to heavy-tailed and normal reward (p < 0.001 and p < 0.01, Benjamini-Hochberg corrected Wilcoxon signed-rank test on non-normalized BICs). **B.** RMSE distribution (median and 50% confidence interval) between observed and bootstrapped learning trajectories for all 5 shape parameter values, *β* ∈ {1,2,4,6,8}. The distributions are sorted for each experimental bird from the best model to the left (smallest RMSE) to the worst model to the right. Note that the best two (three) models are light-tailed 80% (73%) of the time, more than the 60% expected by chance. Besides, in 6 syllables, the best and second-best models are light-tailed, twice as expected by chance, and in 3 syllables, the three-best models are all light-tailed, three times as expected by chance. These are further indications supporting Figures 1G and 1I that the imitation of a single syllable is driven by a light-tailed intrinsic reward. **C.** Similar to Figure 1G with the value of *β* fixed but learning rate *α* inferred with maximum likelihood. **D.** Root-mean-square error (RMSE) between observed and bootstrapped learning trajectories (dashed lines connect same-bird data) comparing the full model where both *α* and *β* were inferred to simpler models where either *β* alone or both *α* and *β* were fixed. The latter, simpler model improves goodness-of-fit (p < 0.05, Benjamini-Hochberg corrected Wilcoxon signed-rank test) and is therefore used in all later simulations. **E.** Left, performance frequency of bigrams containing a shifted song syllable and other motif syllables (% of total bigrams) at experimental start point and end point across birds. Right, same as left for a call. In 5 out of 6 cases, call-containing bigrams did not change their performance frequencies. **F.** Pitch variance analysis justifying the inclusion of daily variance values as additional model parameters: Pitch variance changes over days, decreasing significantly within the last 3 days (at T; orange), compared to the first 3 days (at S; blue) of the experiment (p < 0.001, Benjamini-Hochberg corrected Wilcoxon signed-rank test). However, it remains stable within the same day (early vs. middle vs. late; Benjamini-Hochberg corrected Wilcoxon signed-rank test p > 0.05). Pitch variance increases significantly at the day where the largest pitch shift occurs (learning; purple) compared to two days before and after (p < 0.05, Benjamini-Hochberg corrected Wilcoxon signed-rank test). This may point to a pitch exploration strategy adapted by juvenile birds to learn the new target.

**Figure S3:**
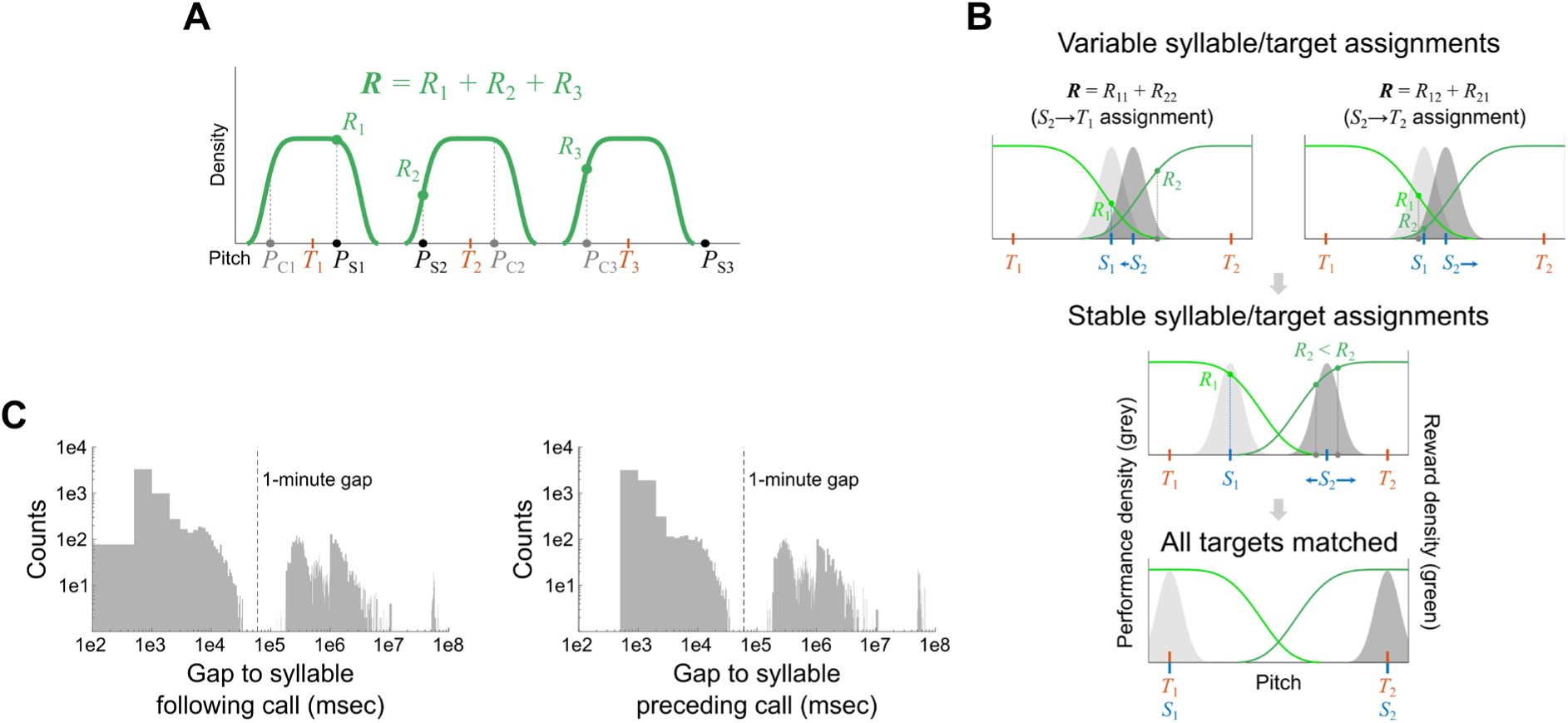
**A.** A hierarchical sum-max reward computation based on light-tailed primitives. A hypothetical example with three target syllables (*T*_1_, *T*_2_, *T*_3_), three song syllables (*S*_1_, *S*_2_, *S*_3_), and three calls (*C*_1_, *C*_2_, *C*_3_) in a bird’s vocal repertoire (*P*; pitch). The global reward *R* is a sum over partial rewards (*R*_1_, *R*_2_, *R*_3_), computed with respect to the most similar syllable within the reward range of each target. If no syllable is within reward range (as is the case with *T*_3_) the partial reward is computed with respect to the most similar call. **B.** Dynamic syllable-target assignments. A hypothetical situation demonstrating dynamic syllable-target assignments during syllable inventory learning: top, two agents (*S*_1_ and *S*_2_) with overlapping performance distributions (light and dark grey) are within the reward range of two target syllables (*T*_1_ and *T*_2_, light and dark green). Assignments between syllables and targets are dynamically determined by the syllable instance just performed (grey circle), and the performance-distribution mean of the other syllable *S*_1_. For example, the *S*_2_ instance depicted on the top left panel would lead to *S*_1_ determining the partial reward associated with target *T*_1_ (*R*_1_ = *R*_11_); and *S*_2_ determining the partial reward associated with target *T*_2_ (*R*_2_ = *R*_22_), which in conjunction with Eq. 10 result in an update of *S*_2_ towards *T*_1_ (*S*_2_ → *T*_1_ assignment); the *S*_2_ instance depicted on the right panel will lead to opposite syllable-target assignment (*S*_2_ → *T*_2_). Since *S*_2_ instances that lead to *S*_2_ → *T*_2_ assignment and, similarly, *S*_1_ instances that lead to *S*_1_ → *T*_1_ assignment will result in a larger inventory reward, and larger mean updates (blue arrows) than instances that lead to *S*_1_ → *T*_2_ and *S*_2_ → *T*_1_ assignments, this dynamical process will lead to a consistent shift of *S*_1_ towards *T*_1_, and of *S*_2_ towards *T*_2_ (middle), and eventually to a match of the complete target vocabulary (bottom). See Figure 2. **C.** Distributions of call-song intervals for bird 3 demonstrating that the average duration of a singing-and-calling period is roughly 1 minute.

**Figure S4.**
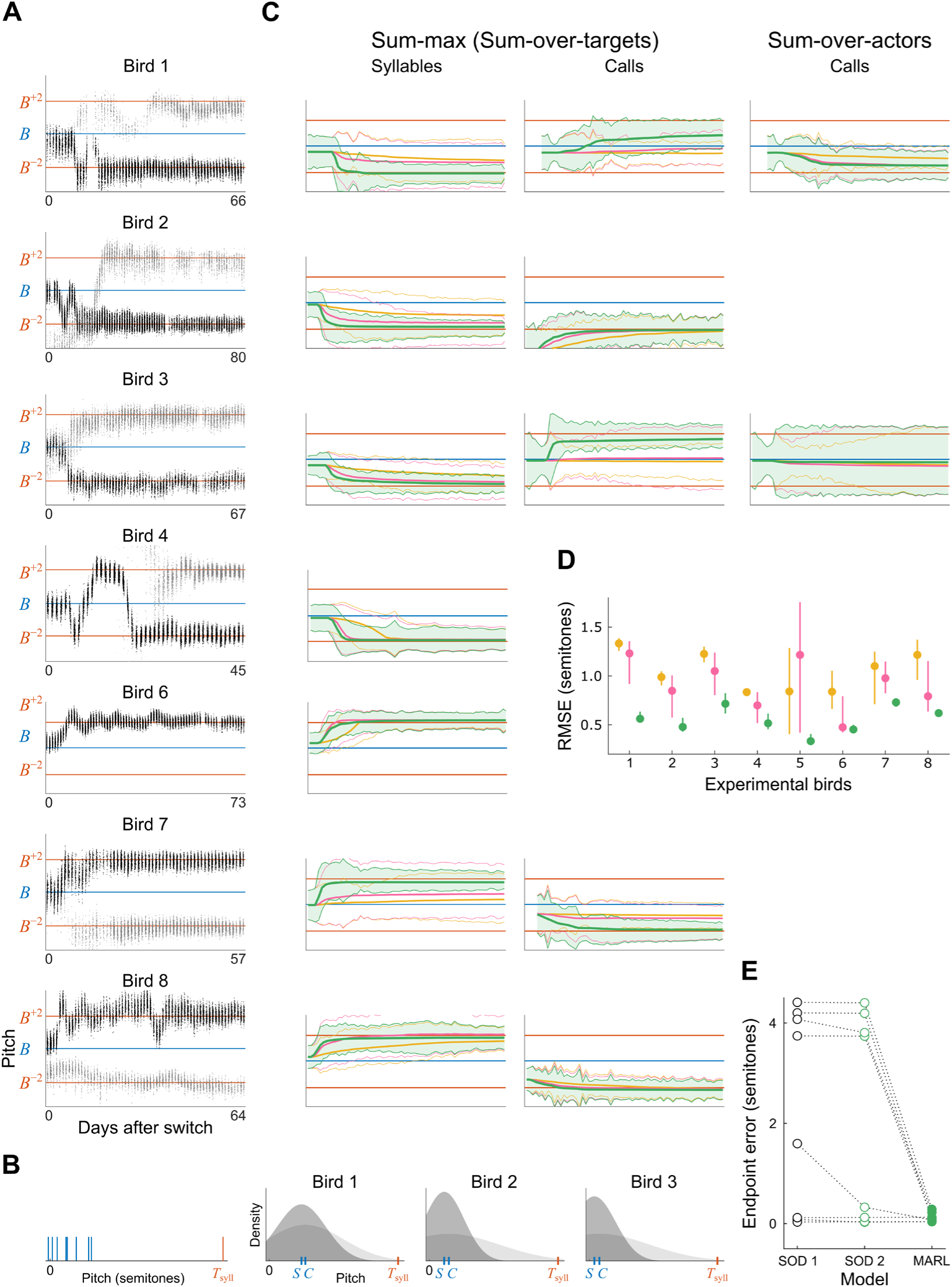
**A.** Observed pitch trajectories for all birds except bird 5 (shown in Figure 2G) that were tutored on an imitation task requiring the matching of two target syllables *B*^+2^ and *B*^−2^ (notation as in Figure 2D, top). ***C.*** Left, when induced to learn two new targets, 7 out of 8 birds shifted a song syllable towards the target *T_syll_* it was closer to at the first day after shift (blue lines). Right, in 3 out of 3 birds where both a syllable *S* and a call *C* means (blue lines) where nearer the same target but the call was closer, the syllable always won – the syllable matched its closest target, even when a call was closer to that target. Performance distributions are shown in gray (dark for *S* and light for *C*). **C.** Bootstrapped syllable (left) and call (middle) pitch trajectories for the experimental birds shown in **A** using the sum-max (i.e., sum-over-targets) MARL model. Note that in bird 2, while the call is assigned to the target syllable *B*^+2^, simulations violated musical chairs, since the call is outside the reward range of that target. Right, bootstrapped call pitch trajectories for birds 1 and 3 using the sum-over-actors MARL model. In these two birds, both the syllable and the call are closer to the same target syllable *B*^−2^, resulting in violation of musical chairs not seen in the data and the sum-max MARL simulations. **D.** RMSE distribution (median and 50% confidence interval) between observed and bootstrapped learning trajectories for heavy-tailed (yellow), normal (magenta), and light-tailed (green) reward distributions, for each experimental bird. **E.** Comparison in the endpoint simulation error between the sum-max MARL model with light-tailed reward (filled green circles) and two sequential-order-dependent (SOD) models, which follows the song learning approach proposed by Fiete and colleagues^8^. SOD 1 (open black circles) uses a long-range binarized reward, while SOD 2 (open green circles) uses the same light-tailed reward distributions as in the sum-max MARL model.

**Figure S5.**
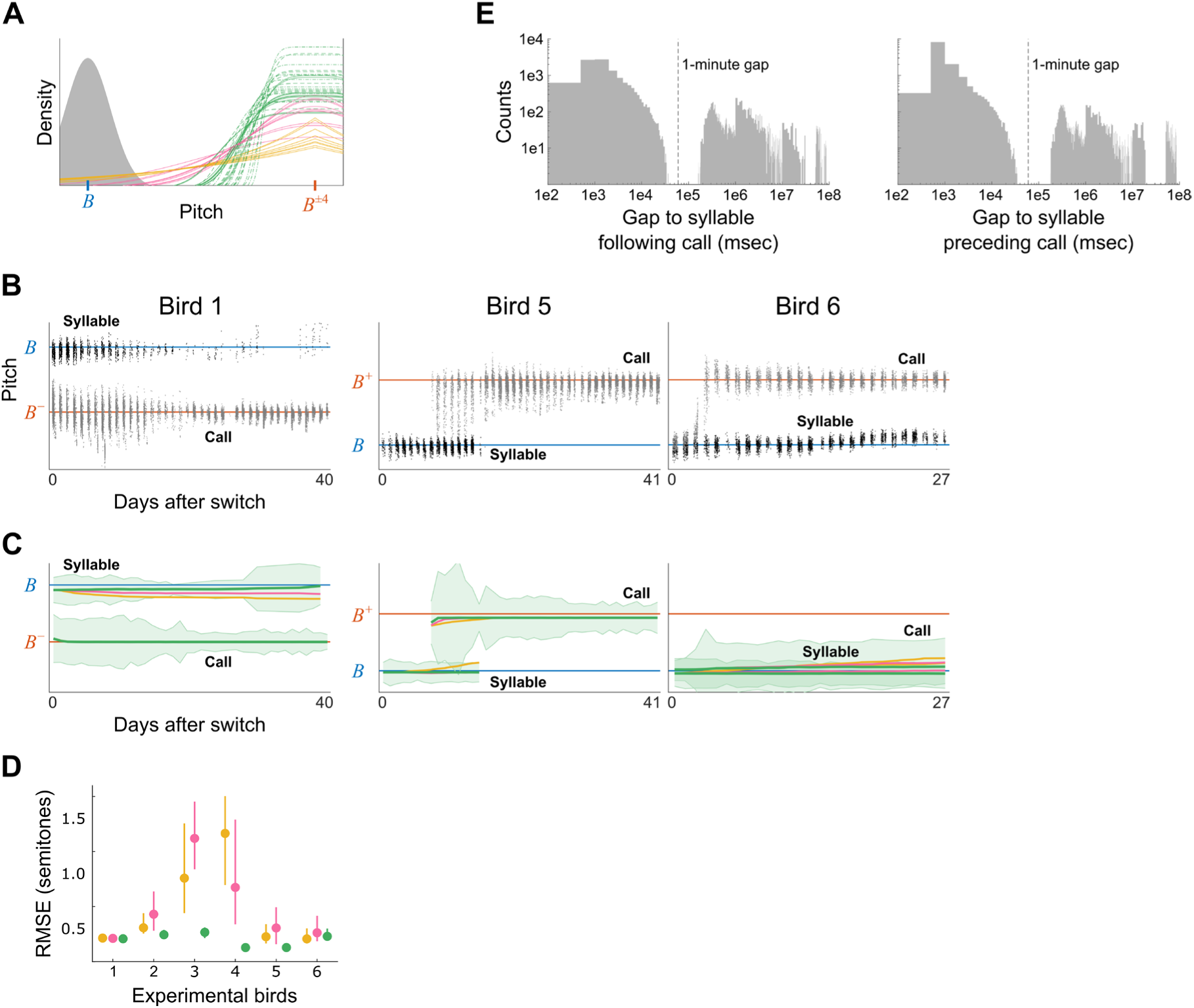
**A.** Reward distributions as estimated in Figure 1 and used in all subsequent simulations (n = 10 for each *β* value; solid green: *β* = 4, dashed green: *β* = 6, dotted green: *β* = 8). Grey, typical performance distribution of syllable B. For the *B*^±4^ target, performance density is outside the light-tailed reward range but within the normal and heavy-tailed reward range (i.e., performance density overlaps with the normal and heavy-tailed reward distributions, but not with the light-tailed distributions). **B-C.** Observed (**B**) and bootstrapped (**C**) syllable and call pitch trajectories for 3 birds tutored on an imitation task requiring the matching of one target syllable that was shifted up or down by 4 semitones. Additional 3 birds are show in Figure 3. **D.** RMSE distribution (median and 50% confidence interval) between observed and bootstrapped learning trajectories for heavy-tailed, normal, and light-tailed reward distributions, for each experimental bird. **E.** Distributions of call-song intervals for bird 3 demonstrating that the average duration of a singing-and-calling period is roughly 1 minute.

## Footnotes

1 We could define 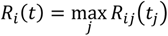 is the partial reward associated with target *T_i_* and *t_i_* ≤ *t* is the last time at which actor *j* produced a vocalization.

